# Ecological assembly dynamics of the seed-borne microbiome in cultivated and wild wheat

**DOI:** 10.1101/2024.12.27.630520

**Authors:** Ezgi Özkurt, Zaywa Mariush, Doreen Landermann-Habetha, Eva H. Stukenbrock

## Abstract

**Background:** Plants host diverse microbial communities that can significantly impact plant health. A portion of these communities is vertically transmitted to the next generation through seeds, influencing early plant development. While seed microbiota composition is known to be shaped by environmental, genetic, and stochastic factors, detailed insights into the factors that govern variation within and between plants of the same species remain limited. Furthermore, the dynamic of the vertically-transmitted microbes and their mode of host colonization are poorly understood.

**Results:** In this study, we characterized the bacterial communities of multiple seeds originating from the same spike of adult wheat plants and demonstrated a high extent of homogeneity in the composition of microorganisms. We propagated wheat seedlings under controlled conditions to monitor host colonization by these microorganisms. Intriguingly, in spite of a relatively homogeneous composition of bacteria in individual seeds, the microbes colonizing early-stage seedlings, particularly during their initial leaf development, vary extensively, even among seedlings of seeds from the same spike. Applying a neutral assembly model, we show that the transmission of microbiomes from seeds to seedlings is primarily governed by selective processes, whereas post-transmission microbial assembly in seedlings is governed by stochastic events.

**Conclusions:** Overall, this study introduces a novel experimental approach for investigating microbiome inheritance in plants. By leveraging high replication of seeds from the same wheat plant and field, we provide new insights into the stochastic nature of seed-associated microbiome composition and transmission in wheat.

## Background

Plant seeds are not standalone entities, but harbor a diversity of endophytes which are transmitted across plant generations (Soluch et al. 2021); (Özkurt et al. 2020); (Kim et al. 2020); (Wassermann et al. 2019). Seed-borne microbes form intimate relationships with the plant hosts: Being the first colonizers, they have advantage to occupy plant tissues before other microbes that are taken up from the environment. Seed-borne microbes are of special interest since they are putative transmitters of essential and beneficial functions across plant generations (Rezki et al. 2018); (Berg and Raaijmakers 2018). For example, seed-borne endophytes aid in rock weathering and plant growth in cacti, thereby supporting establishment of the young plant in barren deserts (Puente et al. 2009). Seed-borne *Sphingomonas melonis*, was shown to confer resistance against another seed-borne bacterium, the pathogen *Burkholderia plantarii*, by disruption of the *B. plantarii* virulence program (Matsumoto et al. 2021). Controlling and manipulating the seed microbiota could allow us to exploit beneficial functions of seed-borne microbes in the context of crop production. For such applied purposes however, fundamental insights into the factors that determine seed-borne microbial assembly as well as the long-term and short-term microbial community dynamics are needed.

Studies aimed at describing the composition of endophytes in seeds, provide evidence of a host and environment-based selection of microbial members. Research so far states that plant genotype, environment, and stochastic effects shape the recruitment of microbes into host seeds (Wassermann et al. 2019); (Klaedtke et al. 2016); (Rybakova et al. 2017); (Rezki et al. 2018). Also, domestication and breeding have been proposed as important factors that influence the assembly of seed-borne microbiomes (Özkurt et al. 2020); (Rybakova et al. 2017); (Adam et al. 2018). Some studies have pointed to a higher stochasticity in microbiome composition in domesticated plant species compared to wild relatives suggesting a relaxation of selection on traits that define microbial associations (Hassani et al. 2020). Seed-associated microbiomes from plants with the same genotype and originating from the same field showed relatively low variability (Özkurt et al. 2020). However, when the seeds were germinated under axenic conditions, the leaf microbiomes in the axenically-grown plants displayed a higher degree of variation. This observation points to a considerable component of stochasticity in the assembly of the seed-borne microbial community in wheat, however the extent of stochasticity among seeds and among early colonizers have so far never been quantified. A controlled experimental approach can enhance our understanding of the assembly dynamics of the inherited microbes in plants and allows for quantitative and qualitative comparative analyses. Plant seeds serve as ideal systems for studying microbial inheritance in metaorganisms, as they provide replicates from the same and different host individuals, facilitating the replication of transmission events.

This study aimed to investigate the dynamics of microbial inheritance in wheat. In our study, we spanned three previously defined stages of microbial inheritance: (Abdelfattah et al. 2023) (i) seed development (plant-to-seed transmission), (ii) seed dormancy (the period between seed maturation and germination), and (iii) seed-to-seedling transmission, hypothesizing that different ecological processes govern each. To characterize microbial inheritance at different stages of seed and early seedling development, we analyzed and compared the bacterial communities of seeds from the same and different plants in a wheat field, as well as in axenic seedlings during their initial leaf development, grown from the same batch of seeds. Additionally, we explored differences between these dynamics in a domesticated wheat and wild wheat variety. Employing metacommunity theory (Miller, Svanbäck, and Bohannan 2018), we conceptualized seedling microbial communities as “local communities” within a broader “metacommunity” of seed or pooled seedling microbiota. In our first model, the metacommunity was defined by bacteria associated with seeds —as these are the only microbial source for the axenic seedlings in our system. Hereby we were able to investigate the initial assembly of seed-borne bacterial communities in seedlings. In the second model, the metacommunity included bacteria from all seedlings across different growth stages, enabling us to explore the ecological dynamics of the seedling bacterial community over time. Our findings suggest that strong selective pressures drive the initial assembly of seed-borne bacterial communities in seedlings, while stochastic events play a more prominent role in shaping the dynamics of seed-borne bacteria as the seedlings grow.

## Material and Methods

### Seed Collections

Our study built on a collection of seeds of the *Triticum aestivum* spring wheat cultivar Quintus (W.von Borries Eckendorf, Leopoldshöhe, Germany) collected in the field of the experimental farm Hohenschulen of Kiel University, North Germany (N 54 18 36, E 9 58 48) in August 2018.

More precisely, we sampled plant seeds from three different plots (1m2 square, each) from the same and different plants **(Figure 1)**. Therefore, in our collections we included 1) seeds originating from the same plant, 2) seeds originating from different plants but from the same plot and finally 3) seeds from different plants collected at different plots **(Figure 1)**. Plants were treated with different fertilizers and pesticides (e.g. Magnesium oxide, N-Gabe, Agent, Biscaya etc.) during their growth in the conventionally managed field.

**Figure 1:**
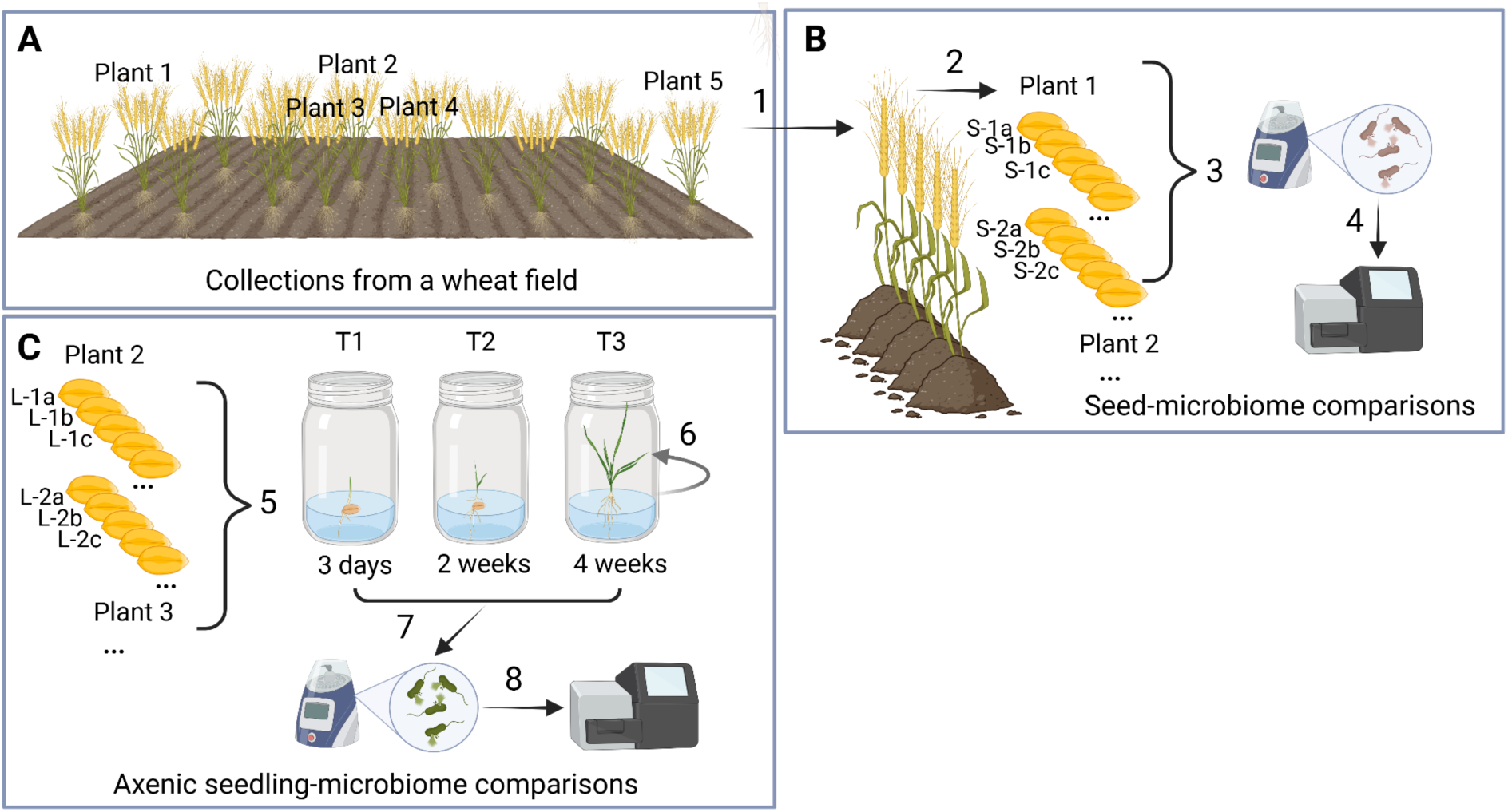
Seed Collections and Experimental Design. **A)** Five adult wheat plants of the cultivar Quintus were collected in a field in August 2018 (1). Two plants were collected from opposite edges of the field-Plant1 and Plant5-, while the other three—Plant2, Plant3, and Plant4—were collected from the center of the field, approximately one meter apart from each other. **B)** Seeds from these plants were harvested (2) (e.g., seeds S-1a, S-1b, and S-1c from Plant 1; seeds S-2a, S-2b, and S-2c from Plant 2), and seed microbiomes within and between individual plants were compared. DNA was extracted from surface-sterilized seeds (3) and processed for 16S rRNA gene profiling (4) to characterize the microbiomes. **C)** To study microbial transmission, a subset of surface-sterilized seeds (seeds from the Plants 2, 3, and 4) was germinated under sterile conditions in jars containing poor nutrient medium (PNM) (5), allowing for the monitoring of vertical microbial transmission from seeds to seedlings (6). Axenic seedlings were harvested at three time points: 3 days (T1), 2 weeks (T2), and 4 weeks (T3). Microbiome composition in seedlings was compared within individual plants and between different plants (e.g., axenic leaves L-1a, L-1b, L-1c from Plant 1; and L-2a, L-2b, L-2c from Plant 2) as well as between different time points (T1, T2, T3) of the seedlings from the same and different plants (7). To do so, the leaf material from seedlings were processed for DNA extraction and 16S rRNA gene sequencing (8). Created in BioRender. Özkurt, E. (2024), https://BioRender.com.

In addition, we included the wild wheat *T. dicoccoides* in our collections to compare the difference in microbiome dynamics between a domesticated wheat species and a wild wheat species. The seed material for these samples were collected in 2006 in South East Turkey (N37 15 33, E37 29 03), which is in the Fertile Crescent region, the center of origin of domesticated bread wheat (Salamini et al. 2002)

### Experimental Design

We aimed to address the ecological processes that shape the seed microbiota in genetically identical plants within a farmer’s field. We examined the relative influence of various factors on microbial community assembly **(Figure 1B)**, focusing on: 1) spike-origin (comparisons based on the different spikes from which the seeds or axenic seedlings originated), 2) growth stage (comparisons of seed and seedling microbiome compositions including different time points of seedling development during axenic propagation), and 3) wheat species (comparisons of seeds and axenic propagated seedlings from the domesticated and wild wheat). Alongside these potential deterministic factors, we also investigated the impact of neutral processes, such as passive dispersal, on the assembly of seed-associated and seed-borne microbiomes.

To assess intra- and inter-plant variation, we compared bacterial communities from seeds of the same spike and seeds from different plants **(Figure 1B, Supp. Table 1**) using 5 to 19 biological replicates of seeds and seedlings across four time points. Only plants with at least five replicates (i.e. five seeds collected from an individual plant) were included in this analysis. Additionally, we applied the Sloan model (Sloan et al. 2006) to predict the stochasticity of seed-associated bacterial communities and the seed-borne bacterial communities in the leaves across different timepoints.

### Processing of seeds for amplicon sequencing

To characterize the composition and dynamics of the seed-associated bacterial communities, we surface-sterilized seeds. The sterilization was mild to eliminate contaminants while preserving microbes from the seed interior and those tightly attached to the surface. Some Quintus wheat seeds were processed for DNA extraction and 16S rRNA sequencing as previously described (Özkurt et al. 2020), while others were used for propagation in closed, sterile jars **(Figure 1B)**.

We propagated surface-sterilized seeds in sterile poor nutrient medium (PNM) medium under 16h light/8h dark cycles at 15°C in a climate chamber (Percival plant growth chambers, CLF PlantClimatics GmbH, Wertingen, Germany). Axenically-grown seedlings were harvested at different stages: T1 (3 days, n=18 from three plants), T2 (two weeks, n=11 from two plants), and T3 (four weeks, n=15 from three plants).

We recorded the seedling development in the closed jars. Three days of growth corresponded to 3-4 cm of leaf growth, two weeks to the emergence of the second leaf, and four weeks to the emergence of the fourth leaf **(Figure 1B)**. Plants with abnormal growth were excluded from the experiment. Leaves were harvested with sterile forceps and DNA was extracted as previously described (Özkurt et al. 2020). Three negative controls were prepared for the sterilization step and for the DNA extraction procedure. These were pooled into one sample and sequenced alongside the plant samples. The negative controls allowed us to ensure the absence of contamination by the handling of the material. DNA from the seed, leaf samples, and negative controls was used to amplify the V5-V7 variable region of the 16S rRNA gene using the 799F/1192R primers (Baker, Smith, and Cowan 2003). Notably, no amplification was observed in the negative controls; therefore, they were excluded from further analyses. Amplicon libraries were paired-end sequenced (2×300 cycles) on an Illumina MiSeq platform (Illumina, San Diego, CA) at the Max Planck Institute for Evolutionary Biology, Plön, Germany.

All samples were randomized during plant growth, DNA extraction, and library preparation.

## Data Analysis

### Pre-processing of the sequencing reads

Reads were pre-processed using the LotuS2 amplicon data analysis pipeline (Özkurt et al. 2022) with default parameters for strict filtering. Filtered reads were clustered into 5,449 OTUs at 97% sequence identity using the UPARSE algorithm (Edgar 2013) integrated within LotuS2. Chimeric sequences were removed *de novo* using VSEARCH (Rognes et al. 2016). A total of 201 OTUs contaminated with PhiX were identified and removed. Additionally, 78% of the remaining OTU sequences showed sequence homology to the *Triticum aestivum* genome (downloaded from EnsemblPlants: https://plants.ensembl.org/Triticum_aestivum/Info/Index) and were filtered out to prevent misclassification of plant-derived reads as bacterial 16S (Bedarf et al. 2021). The remaining 1,156 OTUs were curated using the LULU algorithm (Frøslev et al. 2017) to eliminate erroneous OTUs based on co-occurrence and sequence similarity. After filtering and curation, 236 OTUs (1,807,399 reads) remained in the dataset. These OTUs were taxonomically classified using the LAMBDA3 algorithm (Hauswedell, Singer, and Reinert 2014) based on the SILVA v.138.1 16S/18S rRNA sequence database (Yilmaz et al. 2014). Sequences were aligned using MAFFT (Katoh et al. 2002), and a phylogenetic tree was constructed from this alignment using FASTTREE2 (Price, Dehal, and Arkin 2010).

### Numerical Ecology Analysis

We filtered out OTUs that we could not assign at the domain level and samples with fewer than 1,000 reads. Following these filtering steps, the final OTU matrix comprised 1,715,557 sequence reads representing 95 bacterial OTUs across 76 samples (Supp. Table 2). The OTU table was then rarefied to an even depth to ensure all samples had the same library size (*rarefy_even_depth* function in “phyloseq”) (McMurdie and Holmes 2013) before estimating alpha diversities. Faith’s phylogenetic diversity for each sample and seedling was calculated using the *calculatePD* function in the “biomeUtils” package (S Shetty 2024). Pielou’s evenness for the same samples was calculated using the *evenness* function in the “microbiome” package (Lahti L 2012-2019). Significance of diversity differences between samples was tested with a Wilcoxon test (*stat_compare_means* function in “ggplot2”) (Wilkinson 2011). Alluvial plots of microbiome composition at the class level were created using the *trans_abund* and *plot_bar* functions from the “microeco” package, with integration of the *use_alluvium* function from the “ggalluvial” package (Liu et al. 2021); (Jason Cory Brunson and Quentin D. Read 2023). The OTU table was normalized to relative abundance (*transform_sample_counts* function in “phyloseq”) (McMurdie and Holmes 2013) before calculating beta diversities. A Bray-Curtis metrics was used to estimate between-sample variation in bacterial communities, and the results were visualized in principal coordinate analysis (PCoA) plots using the *get_pcoa* and *ggordpoint* functions in “MicrobiotaProcess” (Xu et al. 2023). The *adonis* function in “vegan” was used to assess the determinants of between-sample variation, while the *betadisper* function calculated community homogeneity for seeds and leaves (Oksanen 2017). The core microbiomes of seeds and seedlings were identified and visualized using the *core* function (detection threshold = 0.0001, prevalence = 0.50) and the *plot_abund_prev* function (mean abundance threshold = 0.01, mean prevalence threshold = 0.99) from the “microbiomeutilities” package (Shetty and Leo Lahti 2020). Finally, differentially abundant OTUs across domesticated wheat timepoints were identified using “MaAsLin2” (Mallick et al. 2021) with thresholds of min_abundance = 0.01, min_prevalence = 0.1, and max_significance = 0.1. Corrected p-values (q-values) for multiple comparisons (Benjamini and Hochberg 1995) were used to assess the significance (q-val < 0.1).

### Inference of the role of neutral processes in microbiome assembly

To assess the role of neutral processes and selection in the assembly of the wheat microbiota, we assessed the fit of the rarefied OTU table of seeds and seedlings to the Sloan Neutral Model (Sloan et al. 2006). This model typically predicts a monotonic relationship where the occurrence frequency of an OTU increases with its mean relative abundance across samples, due to a higher likelihood of dispersal and random sampling from the metacommunity. It reflects the null expectation that OTUs with higher abundance should also appear in more samples. As a result, OTUs located below the predicted trend in the lower-right region of the graph occur in fewer samples than expected based on their average abundance. In contrast, OTUs above the neutral expectation are found in more samples than anticipated. The model also provides the immigration coefficient (*m*), which represents the probability that the loss of a random member in a local community will be replenished through dispersal from the metacommunity. We used the goodness-of-fit (R²) as a quantitative metric to assess the fitness of the data with the neutral model. We used the *fit.scnm* function in the R package “reltools” (Sprockett 2023) and used the approach described by Hassani and co-workers (Hassani et al. 2020) to compute these values and identify bacterial taxa that fit or deviate from the neutral prediction.

For plant microbiota, the definitions of “local community” and “metacommunity” are contextual (Bergmann and Leveau 2022), as the assembly dynamics occur on a micrometer scale. We applied two different interpretations of the Sloan Neutral Model (Sloan et al. 2006), where we defined the metacommunity in two ways: i) first we considered a “constant*m*” by which the immigration rate (*m*) from an external source to the local community is fixed. We applied this by treating pooled seed microbiomes as the “metacommunity” and seedling microbiomes at each time point as “local communities”. ii) secondly, we consider a “dynamic*m*” by which the immigration rate (*m*) varies with the composition of all local communities. Here, we considered pooled seedling microbiomes at each time point (i.e. average of the local communities) as the “metacommunity” and individual seedlings as “local communities”, focusing on transient ecological dynamics such as competition, facilitation, or priority effects during seedling development. Thus, we applied the model as a literal process-based representation (Gotelli and McGill 2006) of community assembly in the first case and as a null model in the second.

## Results

### Sequencing of the seed-borne wheat microbiota

To explore the diversity of bacteria associated with seeds of the wheat cultivar *Quintus*, we amplified and sequenced microbial DNA using the 16S rRNA marker gene. DNA was extracted from individual wheat seeds from different spikes and from the leaves of seedlings propagated under axenic conditions **(Figure 1)**. After filtering of reads and OTUs (see Materials and Methods), our dataset included 76 microbiome samples in total: 20 from seeds and 56 from leaves collected at three seedling stages from four plants in total (Domesticated wheat: T1: n=23, T2: n=13, T3: n=10 & Wild wheat: T1: n=5, T2: n=5) **(Suppl Table 1)**. The final OTU matrix consisted of 1,715,557 sequence reads, representing 95 bacterial OTUs across 76 samples, with 1,054,563 reads derived from seed samples and 660,994 from seedling samples **(Supp. Table 2)**

### Microbial composition in seeds from the same and different wheat plants

We investigated how the seed-borne microbial communities change as bacteria colonize seedlings, and later in the adult plant, the next generation of seeds developing in the spike. We did this by comparing the microbial composition of 1) seeds from the same spike, 2) seeds from spikes of different plants, and 3) seedlings emerging in a sterile environment from surface-sterilized seeds. For the axenically-propagated seedlings, we furthermore included 4) different time points of plant development (five days, two weeks, three weeks). Finally, 5) we compared microbiome dynamics in seedlings from wild and domesticated wheat species.

Seeds were overall predominantly colonized by *Gammaproteobacteria* **(Figure 2A)**. Although the dominant taxa were usually consistent at high taxonomic levels across most seeds, the specific dominant OTUs varied even among seeds from the same wheat spike. For example, in three seeds from the spike of Plant 1, the dominant OTUs were species of *Pantoea, Pseudomonas* and *Uruburuella* respectively. Despite these variations, most taxa (at a minimum relative abundance of 0.0001) were shared among more than 70% of seed samples from the same spike **(Supp. Figure 1A-C)**.

**Figure 2:**
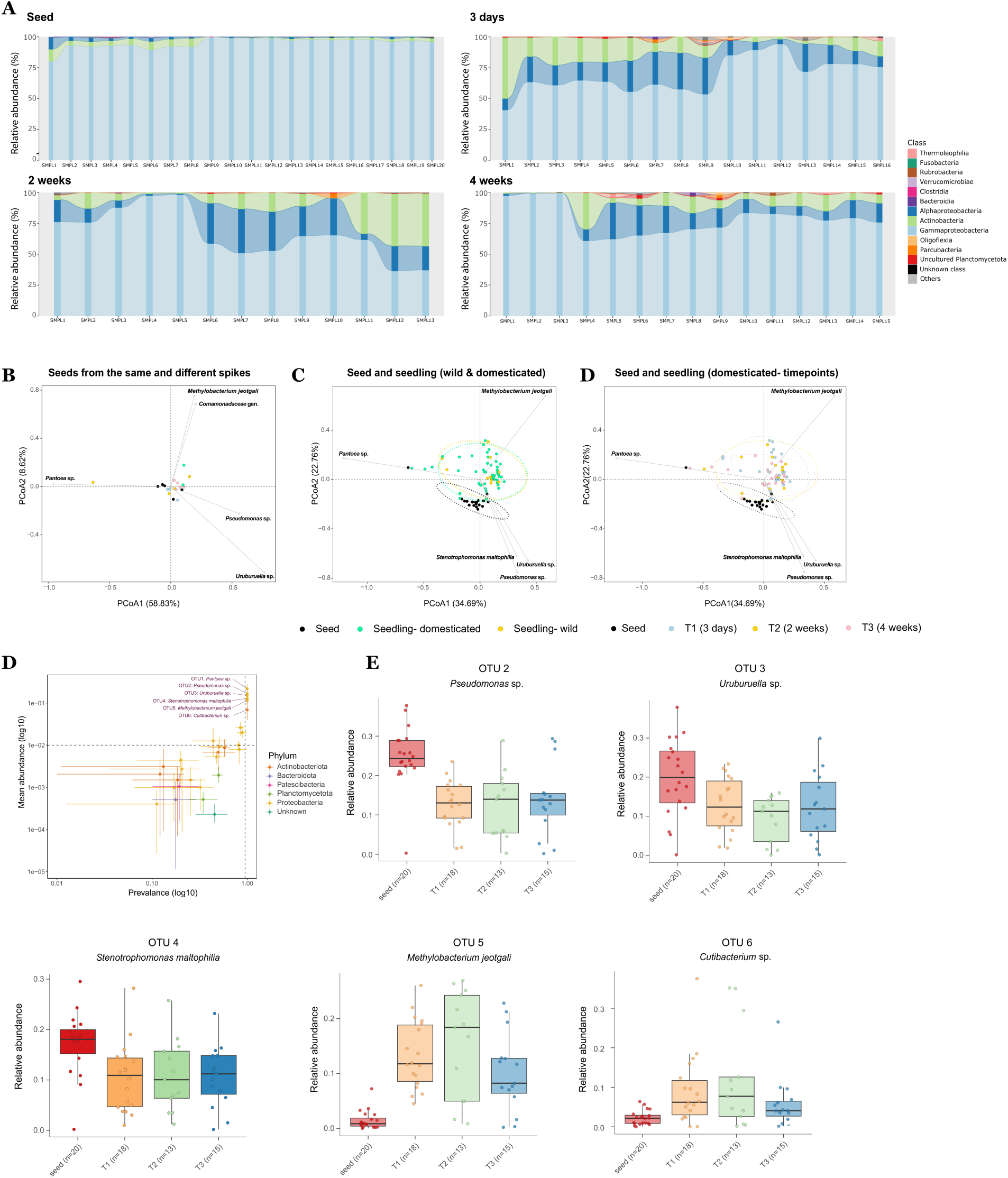
Seed and seedling microbiomes significantly differ in community composition. **A)** Microbiome composition at the class level in seeds and seedlings across different timepoints (3 days, 2 weeks, 4 weeks old plants). Every bar represents one sample. OTUs present in fewer than two samples were excluded from the data used for visualization. **B)** Bray-Curtis distance-based PCoA of bacterial communities in wheat seeds (*T. aestivum*). Each dot represents the bacterial community of an individual seed, with dots of the same color indicating seeds collected from the same wheat spike. **C)** PCoA of bacterial communities in seeds collected from wheat (*T. aestivum*) and seedlings from both wheat and wild wheat (*T. dicoccoides*). Each dot represents the bacterial community of a single seed or seedling from any timepoint. **D)** PCoA of bacterial communities in seeds and seedlings from domesticated wheat at different timepoints (3 days, 2 weeks, 4 weeks). Each dot represents a single seed or seedling. **E)** Core taxa shared between seed and seedlings from wild and domesticated wheat from all time points. These are identified based on prevalence and mean abundance of each OTU in pooled seed and seedling bacterial communities (min abundance = 0.0001 & min prevalence = 50%). Only six OTUs were shared across all samples. **F)** Five differentially abundant OTUs between seeds and seedlings at different timepoints in domesticated wheat. Corrected p-values (q-val) for significant comparisons (qval < 0.1) are shown.

Ordination analysis based on distance matrices showed that microbiome variation among seeds from different spikes (Beta dispersion (e.g. homogeneity) of all the seeds: 0.961) was comparable to the variation within individual spikes (mean of Beta dispersion within individual spikes: 0.1705).

Core taxa shared across all seed samples—including those from the same or different spikes—consisted of *Panteoa* sp., *Pseudomonas* sp., *Uruburuella* sp., *Stenotrophomonas maltophilia*, *Methylobacterium jeotgali*, *Cutibacterium* sp., and *Comamonadaceae* sp **(Supp.** Figure 1D). These core taxa showed little difference between seeds from the same spike and those from different spikes, with the exception that seeds from one spike uniquely included *Tessaracoccus* sp. In their core microbiome **(Supp. Figure 1A-C)**. Overall, microbiome composition showed similar variation within and between wheat spikes, with a consistent core microbiome across seeds.

### Stochastic colonization of seedlings by members of the seed microbiome

We successfully propagated seedlings from the collected seeds of the cultivar Quintus under sterile conditions. We used leaf material of these seedlings to study the fate of seed-borne microorganisms and characterize their distribution in wheat seedlings over four weeks.

As the seeds, also seedlings were predominantly colonized by *Gammaproteobacteria* **(Figure 2A)** however also species of *Alphaproteobacteria* and *Actinobacteria* were abundant in the young plants indicating a shift in taxonomic composition in the seed-to-seedling microbial transmission.

This observation was further supported by an ordination analysis showing that the seedling-associated communities formed a distinct cluster separate from the seed-borne communities (perMANOVA: R² = 14.9%, p < 0.001) **(Figure 2C-D)**. The origin of the seed from the same or different spikes did not explain variation among communities of the seedlings (perMANOVA: p > 0.05). Likewise, we did not detect any effect of the interaction of seedling developmental stage and spike origin (perMANOVA: p > 0.05).

In summary, seed communities, although seeds vary in size and position on the spike, were generally more similar (less dispersed) to each other than the microbial communities found on leaves from the same plant sampled at different time points **(Figure 3B-D)**. This suggests that seeds from the same plant overall comprise a similar composition of microorganisms, however the subsequent colonization of seedlings by the seed-borne microbiome is considerably variable.

**Figure 3:**
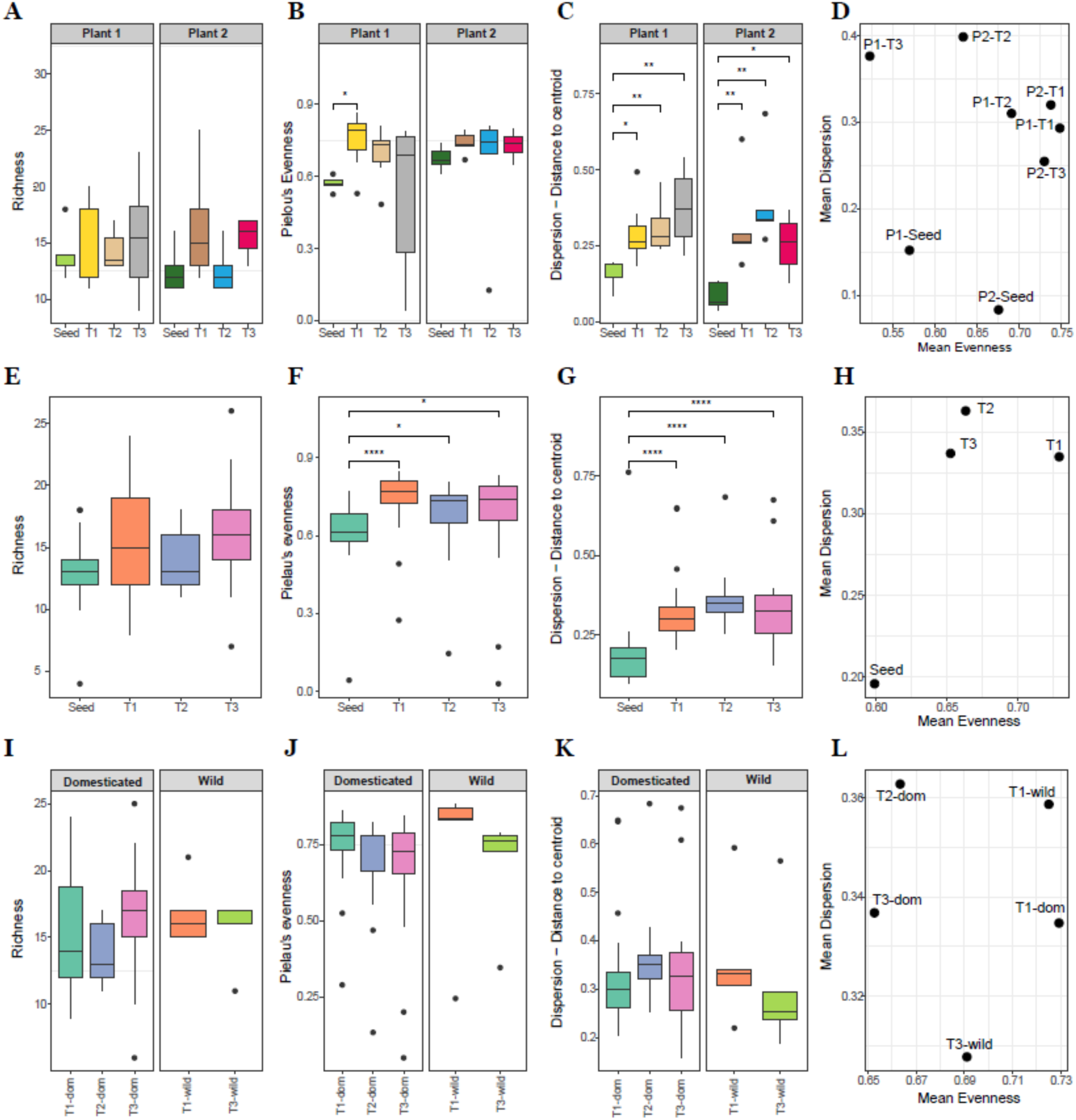
Seed communities are less even but more homogenous than seedling communities, regardless of plant origin and timepoint of the seedlings. A-D) Comparison of diversity (richness), evenness (Pielou’s evenness), and dispersion (distance to centroid in Bray-Curtis ordination plots) between seeds and seedlings from the same wheat plant. **E-H)** Comparison of pooled seeds and seedlings from different plants across timepoints. **I-L)** Comparison of seedlings from different timepoints in domesticated and wild wheat. Significant p-values (pval < 0.05) are indicated in the plots.

We further investigated the specific seedling colonizers based on the OTU classifications. Most taxa were shared among at least 10% of all seed and seedling samples **(Figure 2E)**. Over half (59–64%) of the taxa were shared between seeds and seedlings at each time point for the Quintus wheat, with the highest overlap observed in 4-week-old seedlings (T3) **(Supp. Figure 2A)**. This may be due to late colonizers that increase in abundance after the first two weeks of growth. Only six of the 95 identified taxa were consistently present across all seed and seedling samples: *Pantoea* sp., *Pseudomonas* sp., *Uruburuella* sp., *S. maltophilia*, *M. jeotgali*, and *Cutibacterium* sp. **(Figure 2E)**. The same taxa belonged to the “core taxa” in both seeds and seedlings indicating conserved properties of the seed-borne microbiome composition **(Supp. Figure 3)**. Among these, all, except *Pantoea* sp. were found to exhibit significant differential abundance across different time points in the Quintus wheat **(Figure 2F)**. Of the 95 taxa, 17 met the necessary thresholds for abundance (≥0.0001) and prevalence (≥50%) to be included in the differential abundance analysis, and 11 were identified as significantly differentially abundant (q-value < 0.1) across growth stages **(Supp. Figure 4, Supp. Table 3)**.

In microbiome datasets, rank-abundance curves illustrate the relative abundance of microbial taxa, with the most abundant taxa ranked first, followed by others in descending order. In this study, rank-abundance curves revealed that seeds and seedlings were dominated by a few OTUs, rather than a single taxon, as previously reported in other plants (Chesneau et al. 2022) **(Supp. Figure 5)**. The most dominant OTUs— usually *Pseudomonas* sp. and *Uruburuella* sp. in seeds, along with *Methylobacterium jeotgali* and *Cutibacterium* sp. in three-day-old seedlings—showed a mean relative abundance of 35.2% to 40.7% across both seeds and seedlings. In the Quintus wheat, OTUs displayed a gradual decline in abundance with rank, while in wild wheat seedlings, we observed a more uniform distribution of abundance after rank 1, representing the most dominant taxon. The taxonomy of the dominant OTUs varied, even within seeds and seedlings from the same plant, though one of four OTUs from these taxa consistently dominated the seeds: *Pseudomonas* sp., *Uruburuella* sp., *Pantoea* sp., or *Stenotrophomonas maltophilia*, with the first two being most common (in 6 and 8 out of 20 seeds, respectively). In three-day-old seedlings, these same OTUs were often dominant, along with additional taxa like *M. jeotgali*, *Cutibacterium* sp., and *Comamonadaceae* sp. Notably, although *M. jeotgali* was detected at low abundances in seeds, it frequently emerged as the dominant taxon in three-day-old seedlings (in 6 out of 18 seedlings).

Both *M. jeotgali* and *S. maltophilia*, which were significantly more and less abundant in seedlings than in seeds, respectively, have been previously identified in the seeds of other plants and have been associated with beneficial functions. *Methylobacterium* sp., a common inhabitant of the phyllosphere in various plants (Knief et al. 2010), has been shown to enhance seed germination, improve storability, increase seed vigor through treatments such as seed imbibition and phyllosphere spray inoculation, and protect plant from diseases (Madhaiyan et al. 2015); (Oeum et al. 2024). Similarly, *S. maltophilia* isolated from onion seeds has been reported to possess a broad spectrum of biocontrol and plant growth-promoting characteristics (Guha and Mandal Biswas 2024).

### Dynamics of microbiome composition during early seedling development

We sampled leaves of axenically-grown seedlings at three time points, 3 days, 2 weeks and 3 weeks allowing us to monitor how the microbiome composition changes. In our analysis of the axenic seedling communities, we assessed richness, evenness, and beta dispersion (i.e. homogeneity) among different time points in ordination plots. The measure of evenness, reflecting the distribution of bacterial species within each microbiome, provided us with insights whether communities were dominated by a few species or comprised a more balanced distribution of microbial species. Beta dispersion or homogeneity, calculated based on Bray-Curtis distances from the group centroid in ordination space, measured the homogeneity of group dispersions.

In our closed experimental system, we expected that microbial diversity would not increase in seedlings. As expected, we did not observe significantly higher richness in seeds compared to T1 seedlings (**Figure 3A**). On the other hand, the T1 (3 days old) seedlings in domesticated wheat harbored a significantly more diverse microbial community than seeds, with higher Shannon index values (reflecting greater richness and evenness) and higher Inverse Simpson values (indicating greater evenness) **(Supp. Figure 2B)**. The phylogenetic diversity of seeds and T1 seedlings was however comparable **(Supp. Figure 2B)**. Furthermore, albeit not a significant difference, seedlings from T3 (4 weeks old) exhibited a higher richness compared to T1 and T2 (2 weeks old) seedlings. We speculate that the higher diversity measured in leaves of seedlings is caused by seed-borne microbes of low abundances that are not detected in our analyses of seed-borne microbiome.

Seedling-associated microbiomes, though more variable, exhibited significantly higher evenness compared to seeds, suggesting a compositionally-balanced but variable ecosystem across replicate environments **(Figure 3B-D)**. The findings on the comparison of richness, evenness, and homogeneity of communities between seeds and seedlings remained consistent when we pooled the seeds and seedlings for each time point and conducted a time point-based comparison **(Figure 3E-H)**.

Overall, our study shows that seed-associated microbiomes maintain comparable diversity from seeds to seedlings, suggesting resilience and a strong preservation of a core group of microbes throughout early development, with 59-64% of OTUs shared between seedlings and seeds. Seedlings exhibited higher evenness and dispersion, indicating a more balanced but dynamic microbiome over time.

### Seed-borne microbiome composition is similar in the domesticated and wild wheat seedlings

We previously described differences in the seed-borne microbiome of wild and domesticated wheat species. We here set out to further explore microbiome composition and dynamics of domesticated wheat (*Triticum aestivum*) and a wild relative (*Triticum dicoccoides*). Ordination analysis revealed no distinct clustering of seedling microbiomes according to host species (perMANOVA test, p > 0.05) **(Figure 2C-D)**. Additionally, the interaction between host type and growth stage did not significantly contribute to variation in bacterial community composition among seedlings (perMANOVA test, p > 0.05).

Both domesticated and wild wheat seedlings were predominantly colonized by *Gammaproteobacteria* and *Alphaproteobacteria*, with *Actinobacteria* present to a lesser extent at both growth stages, T1 and T3 **(Figure 2A, Supp. Figure 6)**. However, wild wheat seedlings exhibited lower richness in rare taxa at the class level compared to domesticated wheat seedlings.

Despite these minor differences, wild and domesticated wheat seedlings displayed similar OTU richness, evenness, and homogeneity of microbiome members during early seedling development **(Figure 3I-L)**. Overall, the seed-borne microbiome composition showed no significant variation between domesticated and wild wheat seedlings.

### Selection on vertically-transmitted microbes in seedlings eases during post-germination colonization

The distinct observation of microbiome composition in seeds and seedlings prompt us to further explore the role of selection and stochasticity in microbiome assembly in seeds and leaves. To this end, we first examined whether the initial relative abundance of taxa in seeds could predict their transmission to seedlings. Although not all taxa present in seeds were detected in seedlings **(Supp. Figure 2A)**, we set out to determine if their initial abundance in seeds still played a role in determining transmission success to seedlings. By correlating the relative abundances of taxa in seeds and three-day-old seedlings (T1) from the same wheat plants, we found a strong correlation between seed and seedling microbiomes (Plant 1: R² = 0.61, p < 0.005; Plant 2: R² = 0.52, p < 0.005) **(Figure 4A)**. Rank-abundance curves indicated that each seed was dominated by a few OTUs **(Supp. Figure 5)**, specifically OTU1 (unknown *Pantoea* sp.), OTU2 (unknown *Pseudomonas* sp.), OTU3 (unknown *Uruburuella* sp.), and OTU4 (*Stenotrophomonas maltophilia*), which were also the dominant taxa in the corresponding seedlings. These four OTUs made up nearly 90% of the abundance in seeds, while seedlings had a more even distribution, with five additional taxa contributing to the same total abundance. Overall, our findings suggest that the initial relative abundance of taxa in wheat seeds predicts their transmission to seedlings, but the seedling microbiome is more even than the seed microbiome.

**Figure 4:**
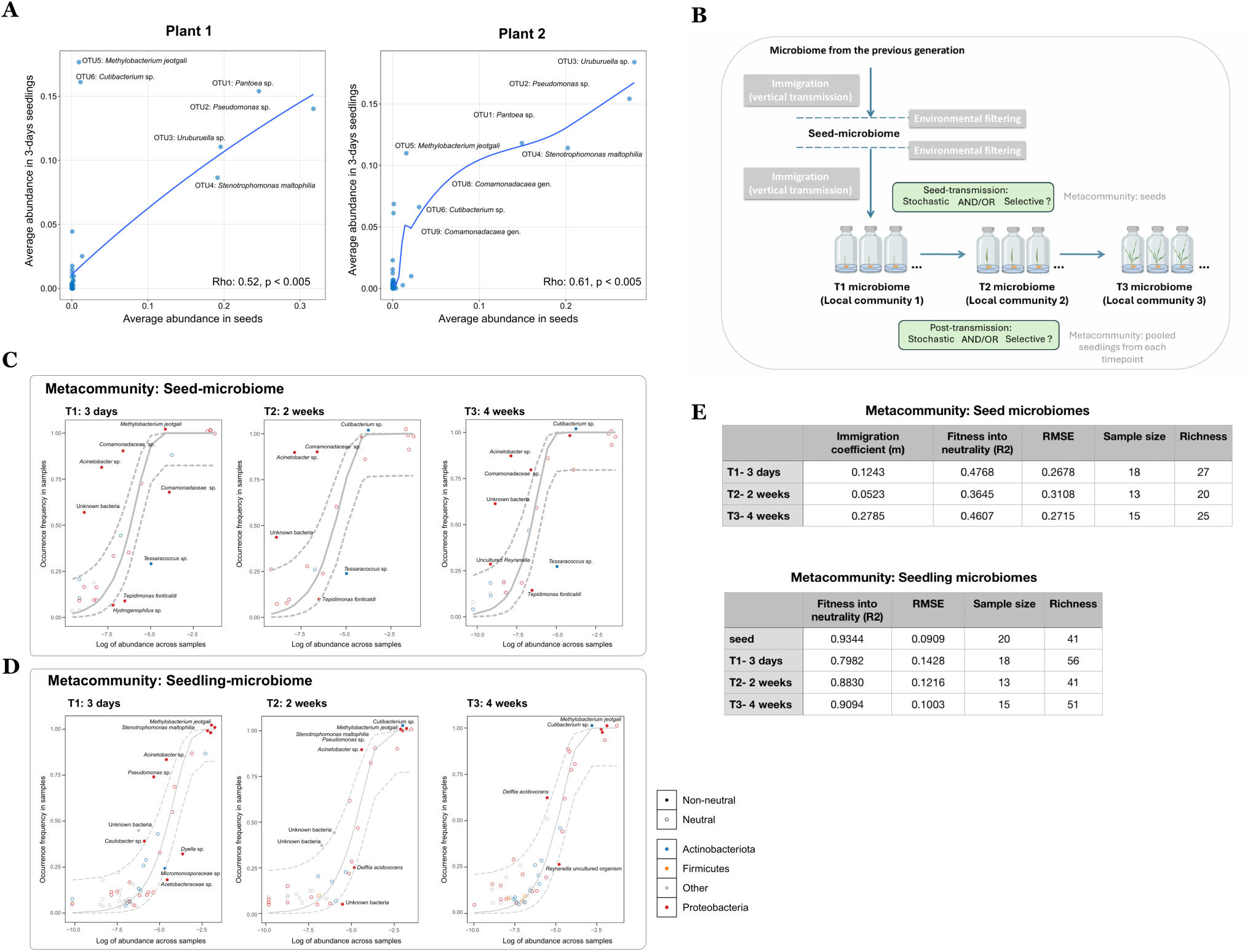
Selection drives vertical microbiome transmission from seeds to seedling leaves, while stochasticity becomes the dominant force shaping seedling microbiomes after transmission. **A)** Comparison of the relative abundance of OTUs in seeds (x-axis) and three-days seedlings (T1) (y-axis). **B)** Conceptual figure depicting ecological forces influencing microbiome assembly in seeds and seedlings, and how they were tested here. **C)** Fit of seedling bacterial communities to the Sloan’s neutral model, with the seed microbiome as the “metacommunity”. Each dot represents an OTU, with the solid line showing the predicted neutral model and dashed lines representing 95% confidence intervals. **D)** Fit of seedling bacterial communities to the Sloan’s neutral model, with the seedlings microbiome from each time point as the “metacommunity”. As in D) each dot represents an OTU, with the solid line showing the predicted neutral model and dashed lines representing 95% confidence intervals. **C-D)** The log of mean relative abundance and detection frequency is plotted to distinguish OTUs that are neutrally distributed (hollow dots) or selected (filled dots). Only OTUs that are identified to be selected by the model were labelled with taxa names. **E)** Summary of the neutrality predictions based on the Sloan’s neutral model.

Next, to estimate the role of neutral processes in the assembly of seed-associated and seed-borne bacterial communities, we evaluated how well the observed taxa distribution fit into the Sloan Neutral Model (Sloan et al. 2006) in both seeds and axenically-grown seedlings (see Material & Methods). Since variations in community composition were not significantly explained by plant origin (intra- and inter-plant comparisons), we combined data from seeds and seedlings into six distinct pools (seeds and seedlings from T1, T2, and T3 from the domesticated wheat and T1 and T3 from the wild wheat). First, we defined the pooled seed microbiomes as the “metacommunity” (i.e., regional pool), from which microbes colonize seedlings, forming “local communities” — the seedling-associated microbiomes **(Figure 4B)**. Our aim was to determine whether the seedling microbiomes reflect a birth-death immigration process, in other words whether chance and immigration are important forces in shaping of seed microbiomes or if selective pressures and competitive dynamics influence this transmission process.

Results showed that the bacterial communities in seedlings significantly deviated from the neutral expectations, with the metacommunity defined as the pooled seed microbiomes (T1: R² = 0.48, T2: R² = 0.36, and T3: R² = 0.46, with R² indicating goodness-of-fit between 0 and 1) **(Figure 4C**, **Figure 4E, Supp. Table 4)**. Notably, the same taxa were mostly categorized within the same prediction class across all three timepoints **(Supp. Figure 7, Supp. Table 5)**. Most taxa that deviated from neutrality, indicating they were under selection pressure, belonged to the *Comamonadaceae* family **(Figure 4C**, **Figure 4E)**. Particularly, *M. jeotgali*, a core member of both seed and seedling microbiomes showed increased abundance in seedlings. This species occurred more frequently than the neutral models predicted, indicating selection for this particular species by the plant **(Figure 4C**, **Figure 4E)**. Finally, the immigration coefficient, which estimates passive microbial dispersal, was higher in seedlings at four weeks (T3: *m* = 0.28) compared to those at three days (T1: *m* = 0.12). Consistent with our finding that bacterial communities in T3 seedlings tend to be more diverse than two weeks old T2 seedlings **(Figure 3A, 3E)**, this finding suggests that late-colonizing microbes may become more abundant and detectable at later stages of seedling development.

Second, we defined the pooled microbiomes at each timepoint as their “metacommunity” (i.e., regional pool), so applied the model as a null hypothesis, and utilized this source pool to evaluate the role of neutral processes in microbiome assembly across these time points **(Figure 4B)**. The seedling microbiome evolves over time, reflecting the dynamics of microbial colonization of newly formed niches, which may be driven by both selective pressures and stochastic processes. Our goal was to estimate the contribution of these processes to the microbial assembly in young seedlings after vertical microbial transmission from seeds.

Bacterial communities in seeds conformed closely to neutral expectations with a relatively high immigration rate (R² = 0.93 & *m* = 0.12) **(Figure 4D**, **Figure 4E, Supp. Table 4).** These conditions describe a community where the random introduction of species through immigration has a stronger influence on community composition than local ecological processes, suggesting a neutral, stochastic assembly process. Therefore, this finding suggests that microbes present in seeds during dormancy are highly explained by a birth-death immigration processSimilarly, communities in seedlings at all three timepoints demonstrated a high fit to neutral expectations (T1: R² = 0.80, T2: R² = 0.88, and T3: R² = 0.91). However, especially in the three-day seedlings (T1), the microbiome community exhibited slightly less stochasticity compared to that in seeds **(Figure 4D**, **Figure 4E)**. Based on the high fitness to neutral expectations and the high variation observed in seedling microbiomes, even among those from the same wheat spike **(Figure 2C-D)**, we conclude that the overall microbiome assembly in seedlings after germination may be driven by stochastic events rather than selective forces. Similarly, the microbial community in wild wheat seedlings showed a strong fit to the neutral model (T1: R² = 0.67; T3: R² = 0.71), though the confidence interval for the fitness was lower since the sample size of the wild wheat was lower than that of the domesticated wheat **(Figure 4E)**.

Our findings suggest that the assembly of vertically-transmitted microbiota in wheat is primarily shaped by selective forces, guiding which microbes successfully establish from seeds to seedlings. In contrast, the microbial dynamics within seedlings, after this initial vertical transmission, are largely governed by neutral processes, introducing a level of stochasticity.

## Discussion

### The core seed-borne microbiome of wheat

In this study, we explored the repeatability of microbial inheritance by comparing the microbiome of seeds collected from the same spike of field-grown wheat plants. Moreover, we tracked microbiome composition through time following the emergence of new leaves in seedlings. Thereby, we demonstrated the assembly of bacterial communities across different seeds of the same plant as well as across seeds from different plants. Moreover, using an axenic system, we characterized the dynamic of bacterial communities from seed to seedling, i.e. in highly distinct ecological niches. Our study is, to our knowledge, the first to present such detailed characterization of wheat seed community assembly and dynamics.

We found that bacterial communities of seeds and axenic seedlings differ significantly in their composition, but this difference was mostly due to changes in the relative abundances of microbes rather than their overall presence or absence. Although there was a significant variation in bacterial composition between seeds and seedlings, we could identify six core bacterial taxa that colonize both seeds and seedlings in all the wheat plants investigated in our study: *Pantoea sp*., *Pseudomonas sp*., *Uruburuella sp., S. maltophilia*, *M. jeotgali*, and *Cutibacterium sp*. Interestingly, we did not observe a significant difference in the composition of bacteria in seedling leaves collected across different timepoints. Similarly, there was no significant difference in the bacterial composition of seedling leaves of the wild wheat relative sampled at different time points suggesting that the seed-borne microbiome that colonizes seedlings is a conserved trait that has not been altered by domestication.

Seeds from the same spike of the wheat cultivar Quintus varied in size according to their position on the spike (data not shown). Yet, we found that the bacterial composition was highly similar, independent of the seed size or position on the spike. In the same way, the bacterial communities of seeds from different plants within the wheat field were highly similar. Notably, wheat seeds were found to be dominated by a few taxa. This observation is different from findings in radish and bean seeds which were found to be dominated by only single taxon (Chesneau et al. 2022). Intriguingly, we found that the taxonomy of the dominant OTUs in the wheat seeds varied, even within seeds or seedlings from the same plant. However, the dominant OTU was always one of the three OTUs assigned to unknown species of the genera *Pantoea*, *Pseudomonas*, and *Uruburuella*. Together, this indicates that replicate seeds or seedlings from the same plant were no more similar to each other than those from different plants.

While some heterogeneity (i.e., variation in the most abundant taxa) was observed in bacterial communities of seeds originating from the same plant, seed-associated microbiomes demonstrated greater similarity compared to seedling microbiomes. Similar heterogeneity in seed microbiomes has also been reported in both individual and pooled samples from rice plants (Kim, Kim, and Lee 2023). However, we did not observe variation in the extent of evenness and dispersion between different plants, suggesting that pooling samples at the plant level, rather than pooling of different plants in a field, may be a more effective approach for studying seed microbiomes. Keeping in mind the challenges to extract microbial DNA from single seeds, we conclude here that several seeds from the same wheat plant could be pooled to study the wheat microbiomes, as also reported for other plant species with larger seeds (Bintarti et al. 2022).

### Seedling-associated microbiomes are less homogeneous than seed-associated microbiomes

While we find highly similar bacterial communities in seeds, we observe a larger variability in bacterial communities in seedlings, even for seedlings that derive from seeds of the same spike. This finding points to a possible role of stochasticity and priority effects in the colonization of young plants by microbes inherited from the mother plant. The observation also demonstrates that the presence of a certain microbial taxa in seeds does not guarantee their transmission to seedlings. While our experiment was not designed to directly track transmission from seed to seedling, our in-depth seed microbiome sequencing still allows us to conduct comparisons of microbial inheritance from seeds to seedlings.

Unlike findings in radish and bean seed-associated microbiomes (Chesneau et al. 2022), our results indicate that the relative abundance of dominant taxa in seeds —representing the initial microbial inoculum of the young plant — can predict the abundance of taxa colonizing axenic seedlings from the same spike. This was especially true for the four most abundant taxa in the wheat seeds, which are species of *Pseudomonas*, *Uruburuella*, *Pantoea*, or the species *Stenotrophomonas maltophilia*. However, there were exceptions; for instance, although *Methylobacterium jeotgali* was detected at low abundances in seeds, it frequently emerged as the dominant taxon in three-day-old seedlings.

When comparing the presence/absence of OTUs between seeds and other timepoints, seeds shared the highest number of OTUs with 4-week-old seedlings (T3). Combined with the higher immigration rate estimated in T3 compared to other timepoints, we speculate that late colonizers established within the plant in detectable amounts over time. However, whether these late colonizers persist when the plant is colonized by environmentally acquired microbes remains a question for future research.

### Deterministic processes govern the vertical transmission of microbes in wheat

In our data analyses, we first treated seed microbiomes as metacommunities and seedling microbiomes as local communities. This allowed us to investigate whether seedling microbiomes are selectively assembled from seed microbiomes or reflect a birth-death immigration process. Our findings indicate that the transmission from seeds to seedlings is predominantly driven by selective processes. This result is expected, given that environmental filtering occurs between seeds and seedlings, two niches with significantly different environmental conditions.

Different tissues have previously been analyzed as metacommunities in a study of stochastic events in the human lung microbiome (Venkataraman et al. 2015). However, to our knowledge, the research presented here is the first to apply such comparative analyses across different tissues and time points in a plant system.

### Vertically-transmitted microbiomes in the seedlings are shaped stochastically over time

We examined whether seedling microbiomes, after vertical transmission, are shaped by selection or stochastic events. Our findings suggest that microbial dynamics in axenically-propagated seedlings over time are primarily driven by stochasticity. Interestingly, the importance of stochastic events is predominant in seedlings of both wild and domesticated wheat. In summary, while a selected portion of the seed microbiome is transmitted to seedlings, stochastic events play a dominant role in the later assembly of axenic seedling microbiomes.

If community assembly was purely governed by neutral processes, the fit of the neutral model would be expected to improve over time (Sieber et al. 2019). Consistent with this expectation, our study shows a gradual increase in the fit of the microbial composition to the neutral model over the course of seedling growth: R² = 0.80 at three days, R² = 0.88 at two weeks, and R² =0.91 at four weeks. These findings further support the role of neutral processes in shaping the vertically-transmitted microbiome community in seedlings.

Microbiome studies have focused on factors that determine the composition of microbial taxa. To this end, a key hypothesis is that the host selects the composition of the microbial taxa (Xiong et al. 2021); (Naylor et al. 2017). This assumption has been challenged in different studies which provide evidence for more stochastic or neutral processes in the assembly of microbial diversity. The role of neutral processes in the assembly of microbiomes in various hosts was investigated by Sieber and co-workers (Sieber et al. 2019). Using a theoretical framework and empirical data, it was possible to show the broad relevance of neutral processes across different model species and tissues. In our study, neutrality estimates for vertical transmission of bacteria from seeds to seedlings (T1: R² = 0.46) and within seedlings across different development stages (T1: R² = 0.80) are comparable to the lowest predicted neutrality in natural *Caenorhabditis elegans* (R² = 0.37) and the highest neutrality in soil compost (R² = 0.87) (Sieber et al. 2019). This highlights a notable shift towards greater stochasticity in the microbiomes after vertical transmission from seed to seedling in wheat.

Overall, this study provides a novel experimental approach for studies of microbiome inheritance in plants and points to the importance of high controlled replication and controlled longitudinal designs.

## Abbreviations

PNM: Poor nutrient medium
T. aestivum: Triticum aestivum
T. dicoccoides: Triticum dicoccoides
OTU: Operational taxonomic unit
R²: Goodness-of-fit

## Acknowledgements

The authors would like to thank M. Amine Hassani for helpful discussions and employees of the experimental farm Hohenschulen of Kiel University for access to wheat fields. The authors thank Víctor M. Flores-Núñez and Román Zapién-Campos for their helpful feedback to a previous version of this manusript.

## Author Contributions

The study was conceived and designed by EHS and EÖ. Domesticated seeds were collected by EÖ and DL. The experiments were performed by EÖ and ZM. Data analysis was conducted by EÖ. The manuscript was written by EÖ and EHS. All authors read and confirmed the manuscript.

## Ethics Declarations and Ethics approval and consent to participate

Not applicable.

## Consent for publication

All authors agree with the publication.

## Supplementary Tables

**Supp. Table 1:** Summary information on the wheat seed collections

**Supp. Table 2:** Summary of the amplicon sequencing data

**Supp. Table 3:** Differentially abundant OTUs identified between seeds and seedlings at different timepoints in domesticated wheat.

**Supp. Table 4:** Summary statistics of the fitness of the bacterial communities into Sloan model in seeds and seedlings. The goodness-of-fit of the Sloan model and immigration coefficient are denoted by R² and *m*, respectively. RMSE indicates the root-mean-square error. AIC.pois and BIC.pois indicate Akaike and Bayesian information criterion for the poisson model, respectively.

**Supp. Table 5:** Prediction class of each bacterial OTU in seeds and seedlings into the neutrality based on the Sloan model.

## Supplementary Figures

**Supp. Figure 1: Core taxa shared among seeds from the same wheat spike (A–C) and core taxa shared between seeds from different wheat spikes (D).** These are identified based on prevalence and mean abundance of each OTU in pooled seed bacterial communities (min abundance = 0.0001 & min prevalence = 50%).

**Supp. Figure 2: A) Percentage of shared OTUs between seeds and seedlings from domesticated wheat.** The black portion of the bars represents the percentage of OTUs shared between seedlings at each time point and seeds**. B) OTU diversity in seeds and seedlings from domesticated and wild wheat.** Each dot represents the diversity within an individual seed or seedling sample. Only significant comparisons are displayed.

**Supp. Figure 3: Core taxa shared between seed and seedlings from domesticated wheat from all time points.** These are identified based on prevalence and mean abundance of each OTU in pooled seed and seedling bacterial communities (min abundance = 0.0001 & min prevalence = 50%). Only six OTUs were shared across all samples.

**Supp. Figure 4: Taxa with significantly different abundances between different time points of samples from domesticated wheat.**

**Supp. Figure 5: Rank-abundance curves (rank1 to rank7) of bacterial OTUs associated with individual seeds and seedlings.** Significance values for p < 0.0005 is denoted by ***, p < 0.005 as **, p < 0.05 as *, and p > 0.05 as “ns”.

**Supp. Figure 6: Microbiome composition at the class level in seedlings across different timepoints (3 days and 4 weeks).** OTUs present in fewer than two samples were excluded from the data used for visualization.

**Supp. Figure 7: PCoA based on Jaccard metrics depicting the composition of neutral and non-neutral partitions of seedling microbiomes, showing high similarity across timepoints.** OTUs are classified into partitions based on prediction classes.

