## Supplemental Figures for "Ecological assembly dynamics of the seed-borne microbiome in cultivated and wild wheat": Supp_Fig4_updated.pdf

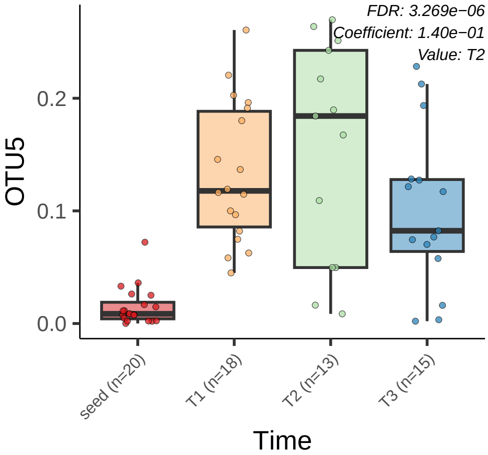

FDR: 3.269e−06
Coefficient: 1.40e−01
Value: T2
OTU5
0.2
0.1
0.0
seed (n=20)
T1 (n=18)
T2 (n=13)
T3 (n=15)
Time

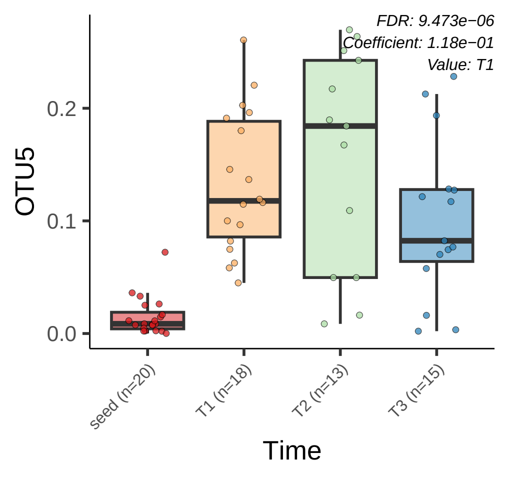
FDR: 9.473e−06
Coefficient: 1.18e−01
Value: T1
OTU5
0.2
0.1
0.0
seed (n=20)
T1 (n=18)
T2 (n=13)
T3 (n=15)
Time

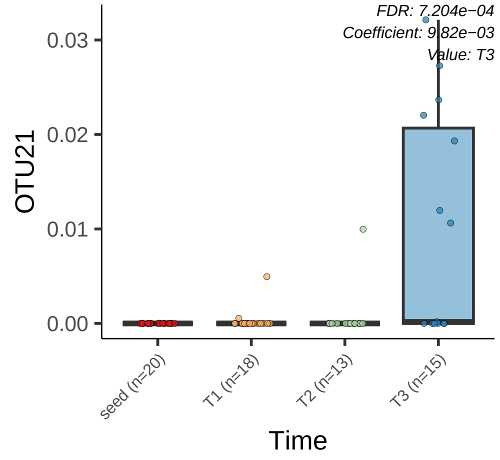
FDR: 7.204e−04
Coefficient: 9.82e−03
Value: T3
OTU21
0.03
0.02
0.01
0.00
seed (n=20)
T1 (n=18)
T2 (n=13)
T3 (n=15)
Time

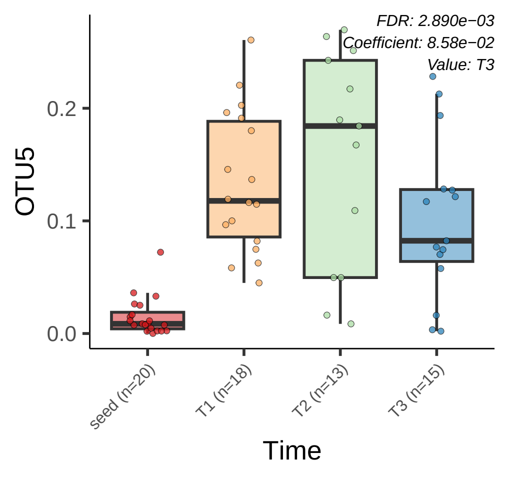
FDR: 2.890e−03
Coefficient: 8.58e−02
Value: T3
OTU5
0.2
0.1
0.0
seed (n=20)
T1 (n=18)
T2 (n=13)
T3 (n=15)
Time

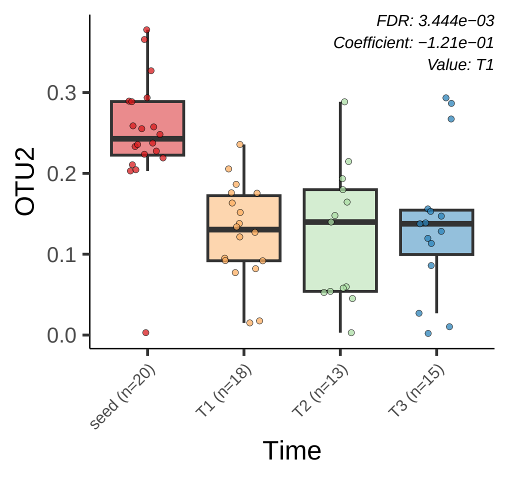

FDR: 3.444e−03
Coefficient: −1.21e−01
Value: T1
OTU2
0.0
0.1
0.2
0.3
seed (n=20)
T1 (n=18)
T2 (n=13)
T3 (n=15)
Time

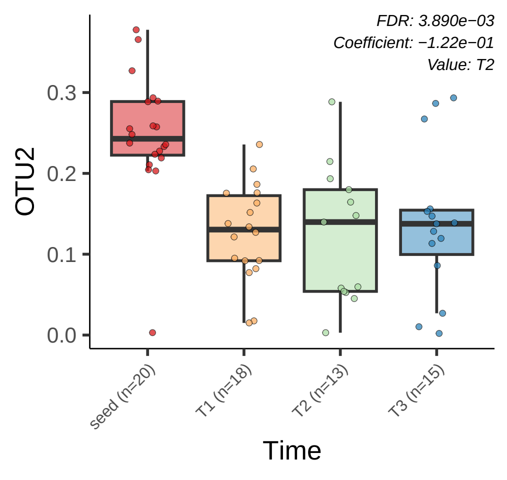

FDR: 3.890e−03
Coefficient: −1.22e−01
Value: T2
OTU2
0.3
0.2
0.1
0.0
seed (n=20)
T1 (n=18)
T2 (n=13)
T3 (n=15)
Time

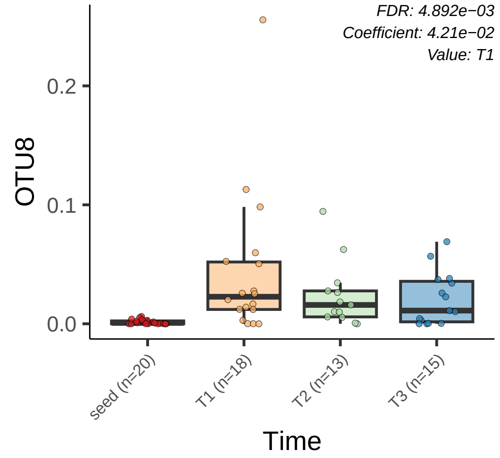

FDR: 4.892e−03
Coefficient: 4.21e−02
Value: T1
OTU8
0.2
0.1
0.0
seed (n=20)
T1 (n=18)
T2 (n=13)
T3 (n=15)
Time

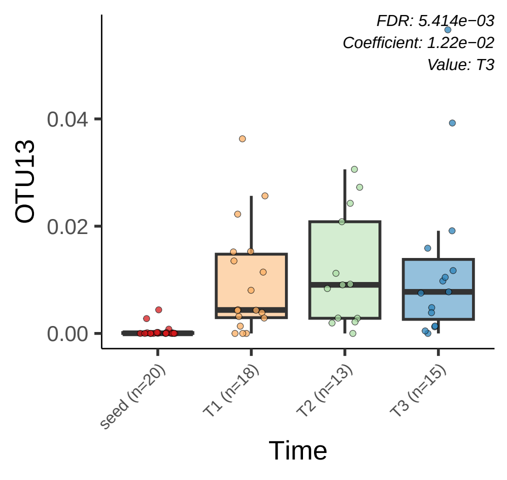
FDR: 5.414e−03
Coefficient: 1.22e−02
Value: T3
OTU13
0.04
0.02
0.00
seed (n=20)
T1 (n=18)
T2 (n=13)
T3 (n=15)
Time

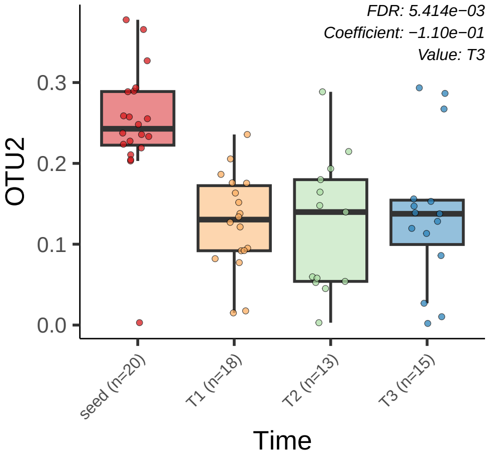

FDR: 5.414e−03
Coefficient: −1.10e−01
Value: T3
OTU2
0.3
0.2
0.1
0.0
seed (n=20)
T1 (n=18)
T2 (n=13)
T3 (n=15)
Time

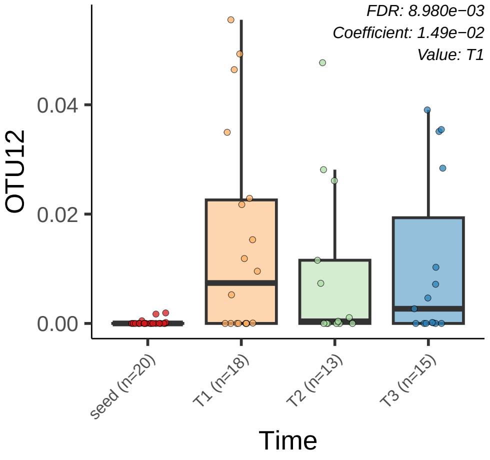

FDR: 8.980e−03
Coefficient: 1.49e−02
Value: T1
OTU12
0.04
0.02
0.00
seed (n=20)
T1 (n=18)
T2 (n=13)
T3 (n=15)
Time

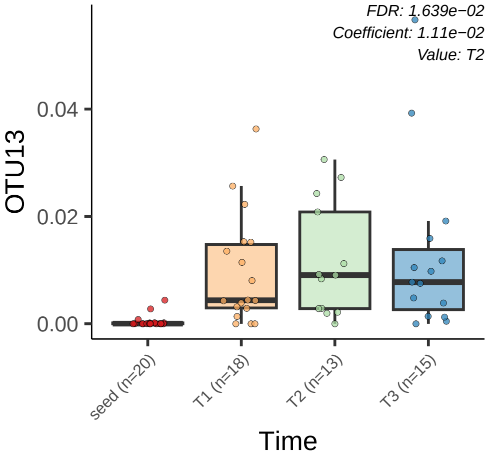
FDR: 1.639e−02
Coefficient: 1.11e−02
Value: T2
0.04
0.02
0.00
OTU13
seed (n=20)
T1 (n=18)
T2 (n=13)
T3 (n=15)
Time

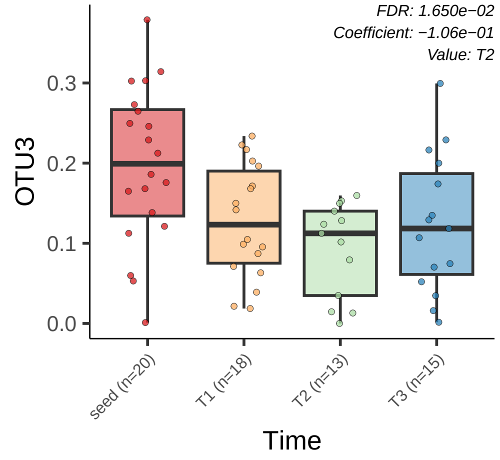
FDR: 1.650e−02
Coefficient: −1.06e−01
Value: T2
OTU3
0.0
0.1
0.2
0.3
seed (n=20)
T1 (n=18)
T2 (n=13)
T3 (n=15)
Time

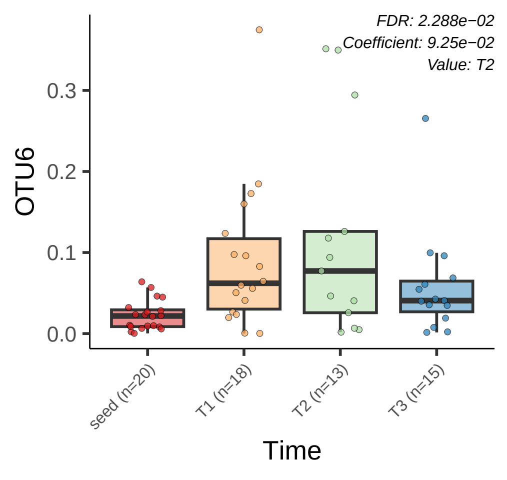
FDR: 2.288e−02
Coefficient: 9.25e−02
Value: T2
OTU6
0.0
0.1
0.2
0.3
seed (n=20)
T1 (n=18)
T2 (n=13)
T3 (n=15)
Time

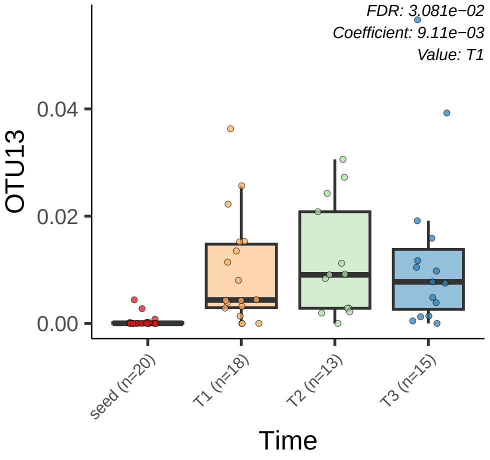

FDR: 3.081e−02
Coefficient: 9.11e−03
Value: T1
OTU13
0.04
0.02
0.00
seed (n=20)
T1 (n=18)
T2 (n=13)
T3 (n=15)
Time

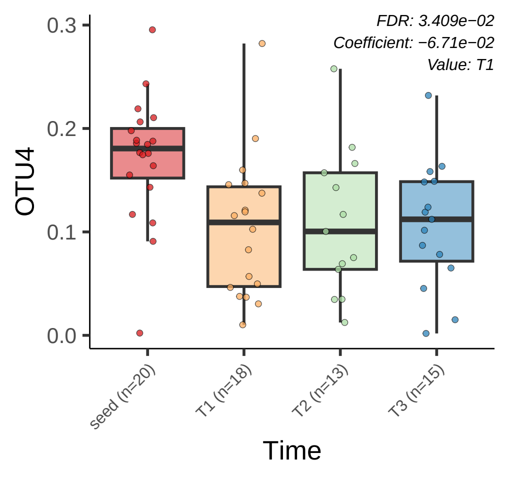
FDR: 3.409e−02
Coefficient: −6.71e−02
Value: T1
OTU4
0.0
0.1
0.2
0.3
seed (n=20)
T1 (n=18)
T2 (n=13)
T3 (n=15)
Time

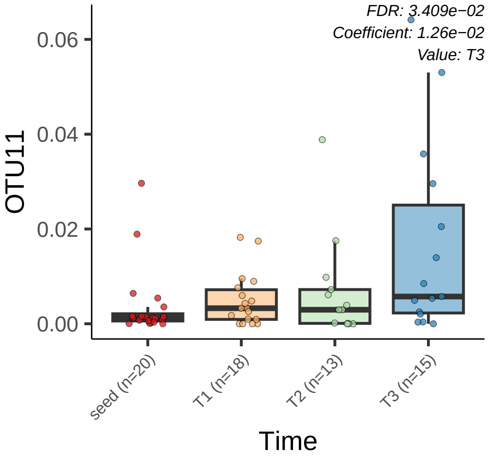

FDR: 3.409e−02
Coefficient: 1.26e−02
Value: T3
OTU11
0.06
0.04
0.02
0.00
seed (n=20)
T1 (n=18)
T2 (n=13)
T3 (n=15)
Time

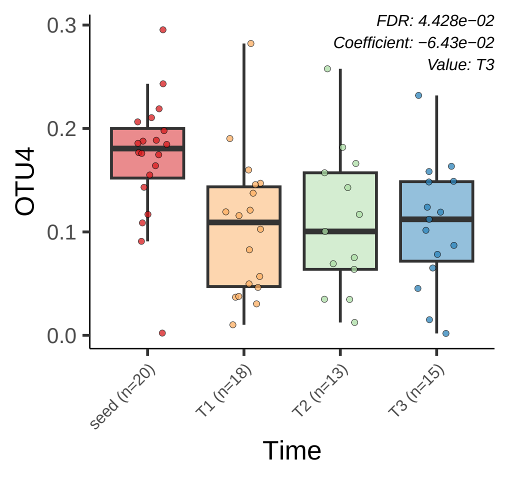
FDR: 4.428e−02
Coefficient: −6.43e−02
Value: T3
OTU4
0.3
0.2
0.1
0.0
seed (n=20)
T1 (n=18)
T2 (n=13)
T3 (n=15)
Time

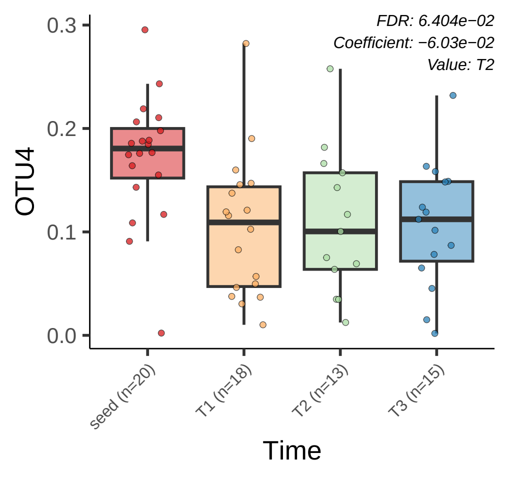
FDR: 6.404e−02
Coefficient: −6.03e−02
Value: T2
0.3
0.2
0.1
0.0
OTU4
seed (n=20)
T1 (n=18)
T2 (n=13)
T3 (n=15)
Time

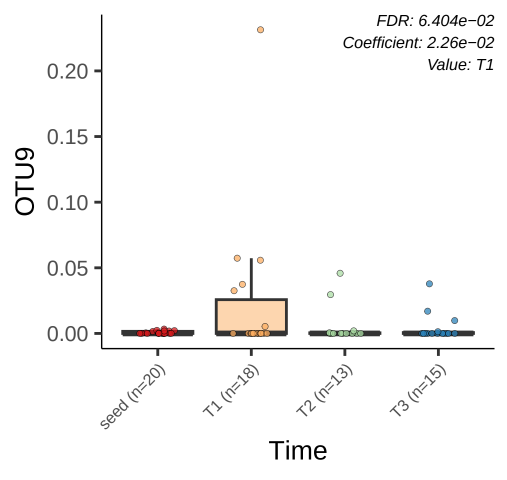
FDR: 6.404e−02
Coefficient: 2.26e−02
Value: T1
0.20
0.15
0.10
0.05
0.00
OTU9
seed (n=20)
T1 (n=18)
T2 (n=13)
T3 (n=15)
Time

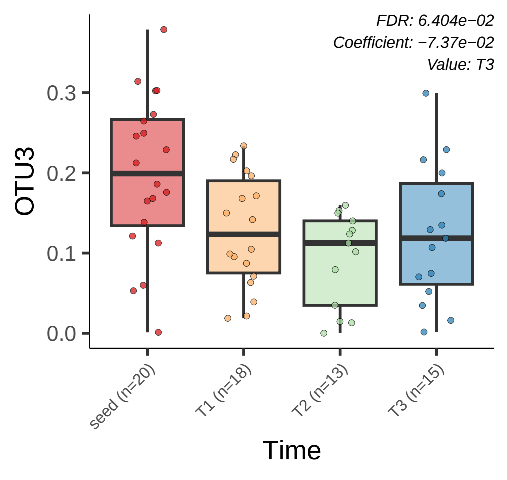
FDR: 6.404e−02
Coefficient: −7.37e−02
Value: T3
OTU3
0.3
0.2
0.1
0.0
seed (n=20)
T1 (n=18)
T2 (n=13)
T3 (n=15)
Time

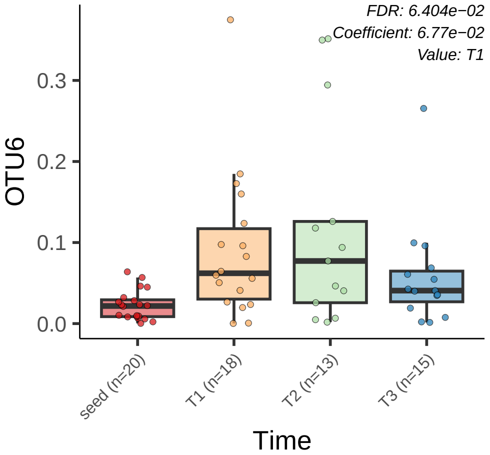
FDR: 6.404e−02
Coefficient: 6.77e−02
Value: T1
OTU6
0.0
0.1
0.2
0.3
seed (n=20)
T1 (n=18)
T2 (n=13)
T3 (n=15)
Time

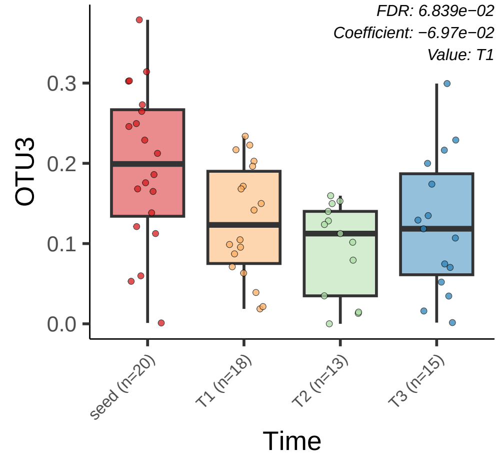
FDR: 6.839e−02
Coefficient: −6.97e−02
Value: T1
OTU3
0.0
0.1
0.2
0.3
seed (n=20)
T1 (n=18)
T2 (n=13)
T3 (n=15)
Time

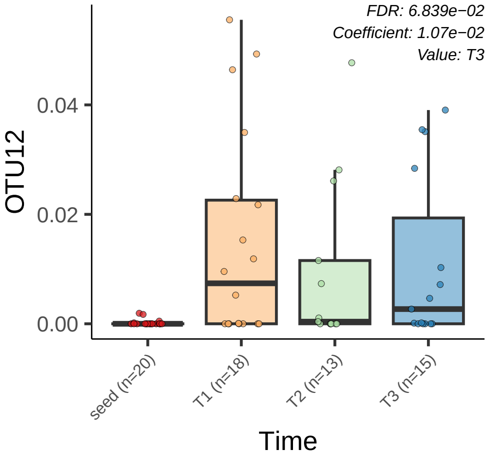
FDR: 6.839e−02
Coefficient: 1.07e−02
Value: T3
0.04
0.02
0.00
OTU12
seed (n=20)
T1 (n=18)
T2 (n=13)
T3 (n=15)
Time

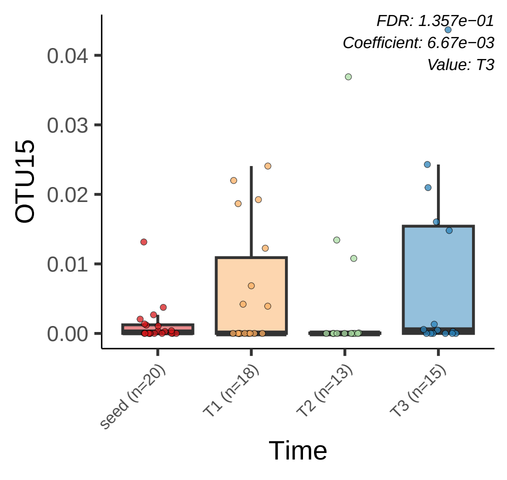
FDR: 1.357e−01
Coefficient: 6.67e−03
Value: T3
0.04
0.03
0.02
0.01
0.00
OTU15
seed (n=20)
T1 (n=18)
T2 (n=13)
T3 (n=15)
Time

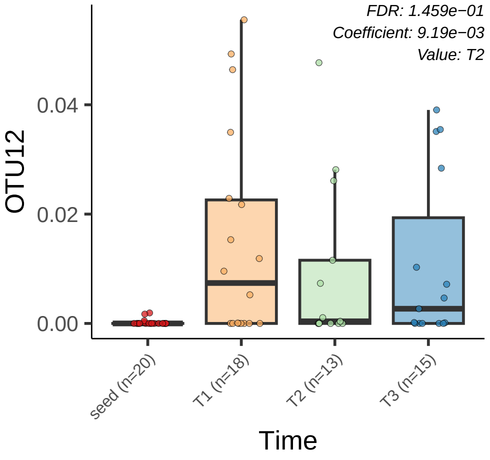

FDR: 1.459e−01
Coefficient: 9.19e−03
Value: T2
OTU12
0.04
0.02
0.00
seed (n=20)
T1 (n=18)
T2 (n=13)
T3 (n=15)
Time

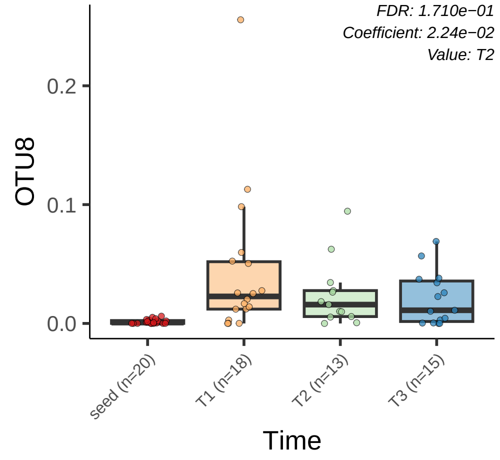

FDR: 1.710e−01
Coefficient: 2.24e−02
Value: T2
OTU8
0.2
0.1
0.0
seed (n=20)
T1 (n=18)
T2 (n=13)
T3 (n=15)
Time

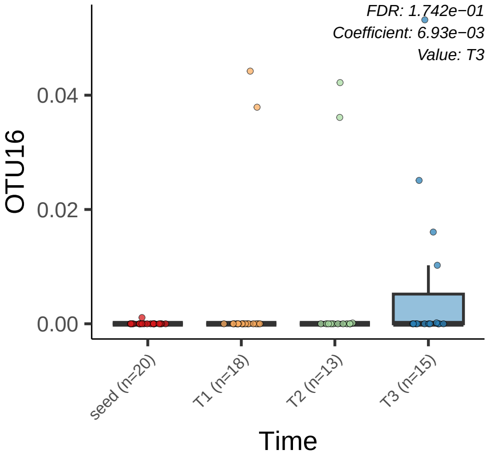
FDR: 1.742e−01
Coefficient: 6.93e−03
Value: T3
OTU16
0.04
0.02
0.00
seed (n=20)
T1 (n=18)
T2 (n=13)
T3 (n=15)
Time

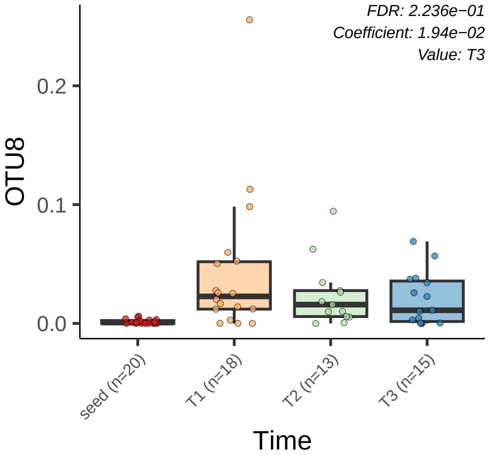

FDR: 2.236e−01
Coefficient: 1.94e−02
Value: T3
OTU8
0.2
0.1
0.0
seed (n=20)
T1 (n=18)
T2 (n=13)
T3 (n=15)
Time

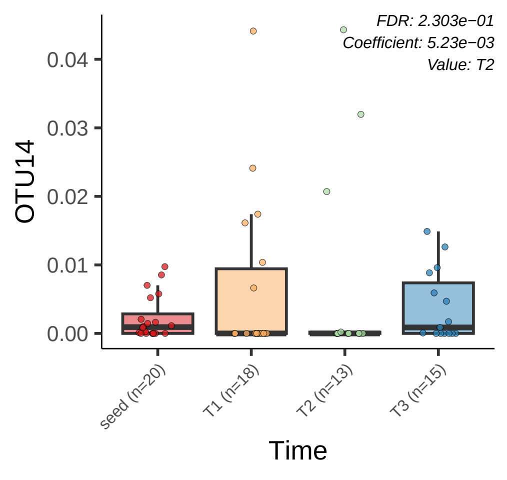
FDR: 2.303e−01
Coefficient: 5.23e−03
Value: T2
OTU14
0.04
0.03
0.02
0.01
0.00
seed (n=20)
T1 (n=18)
T2 (n=13)
T3 (n=15)
Time
