## Supplementary figures and images for "Ecological assembly dynamics of the seed-borne microbiome in cultivated and wild wheat"

### Supp_Fig2_updated.pdf

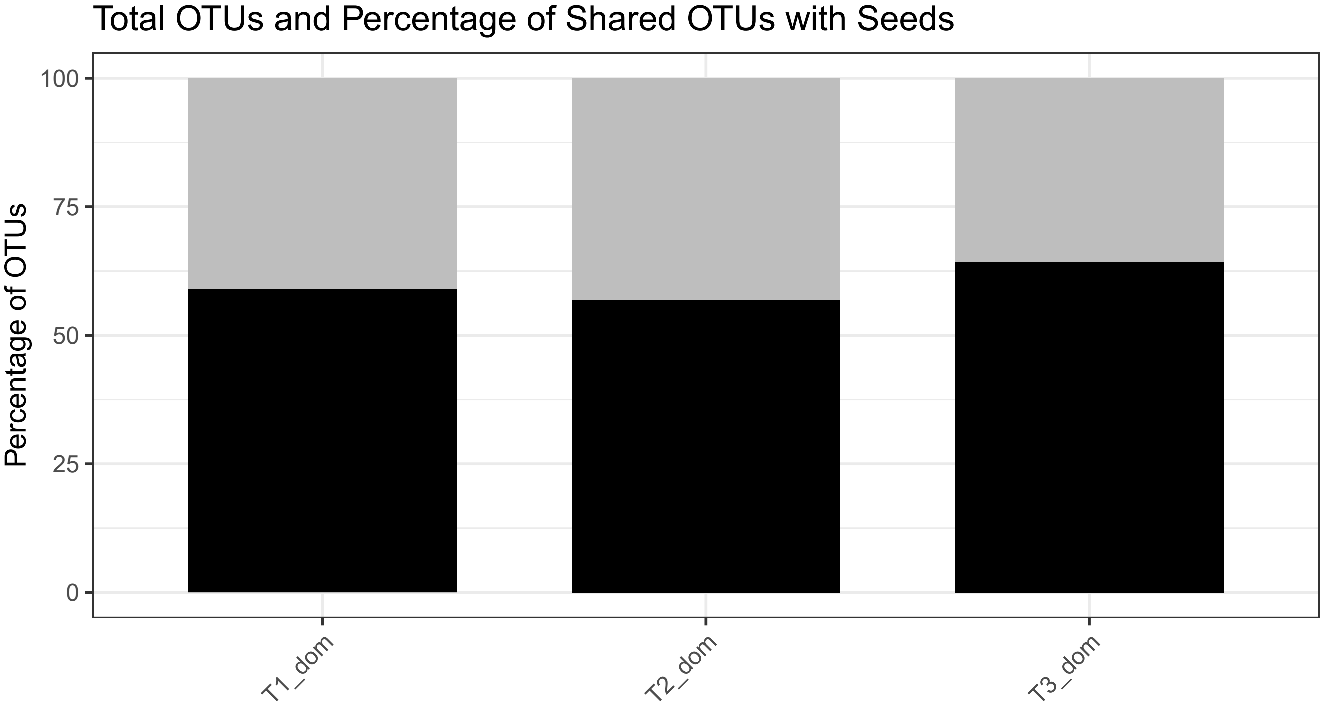

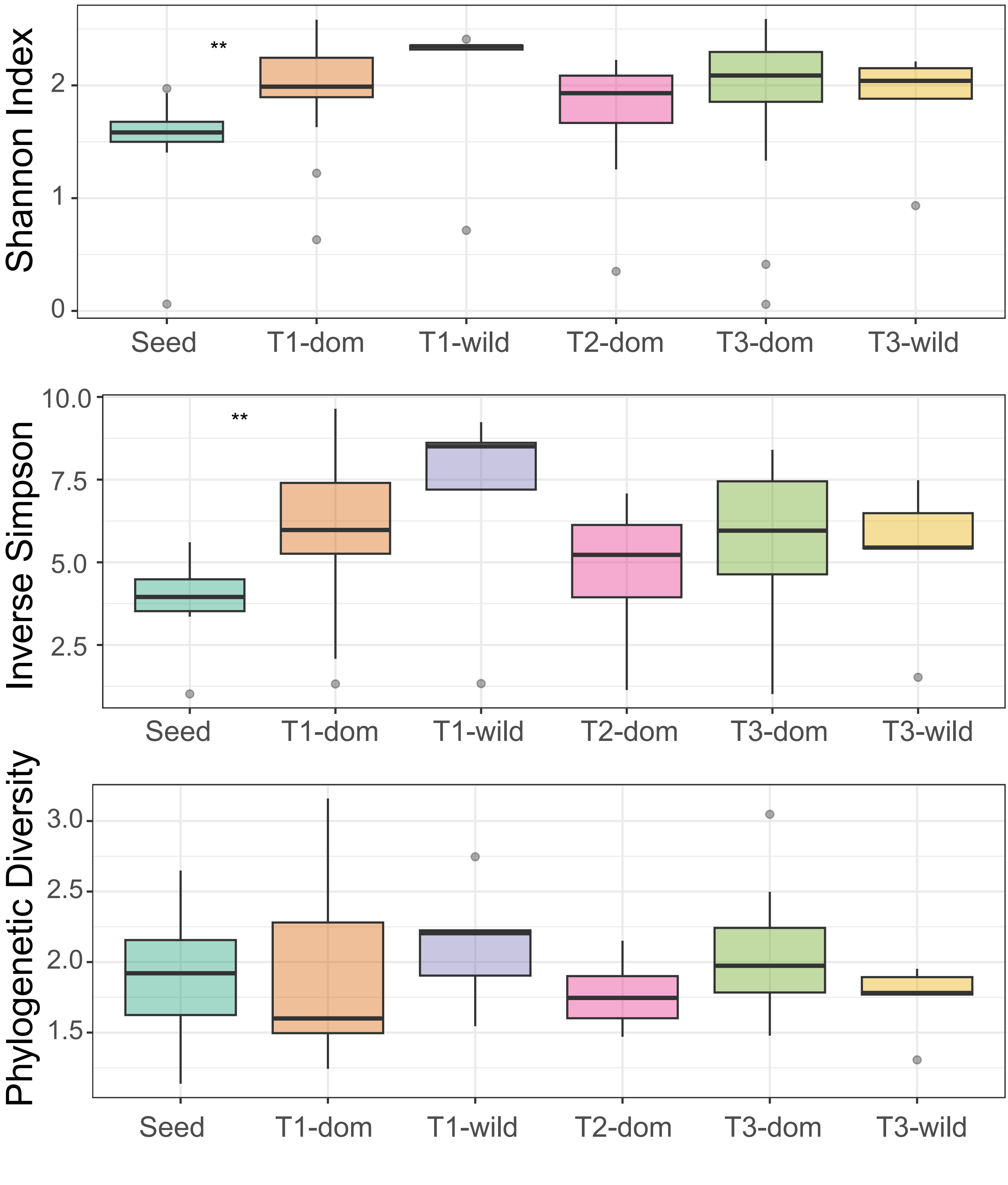

### Supp_Fig3_updated.pdf

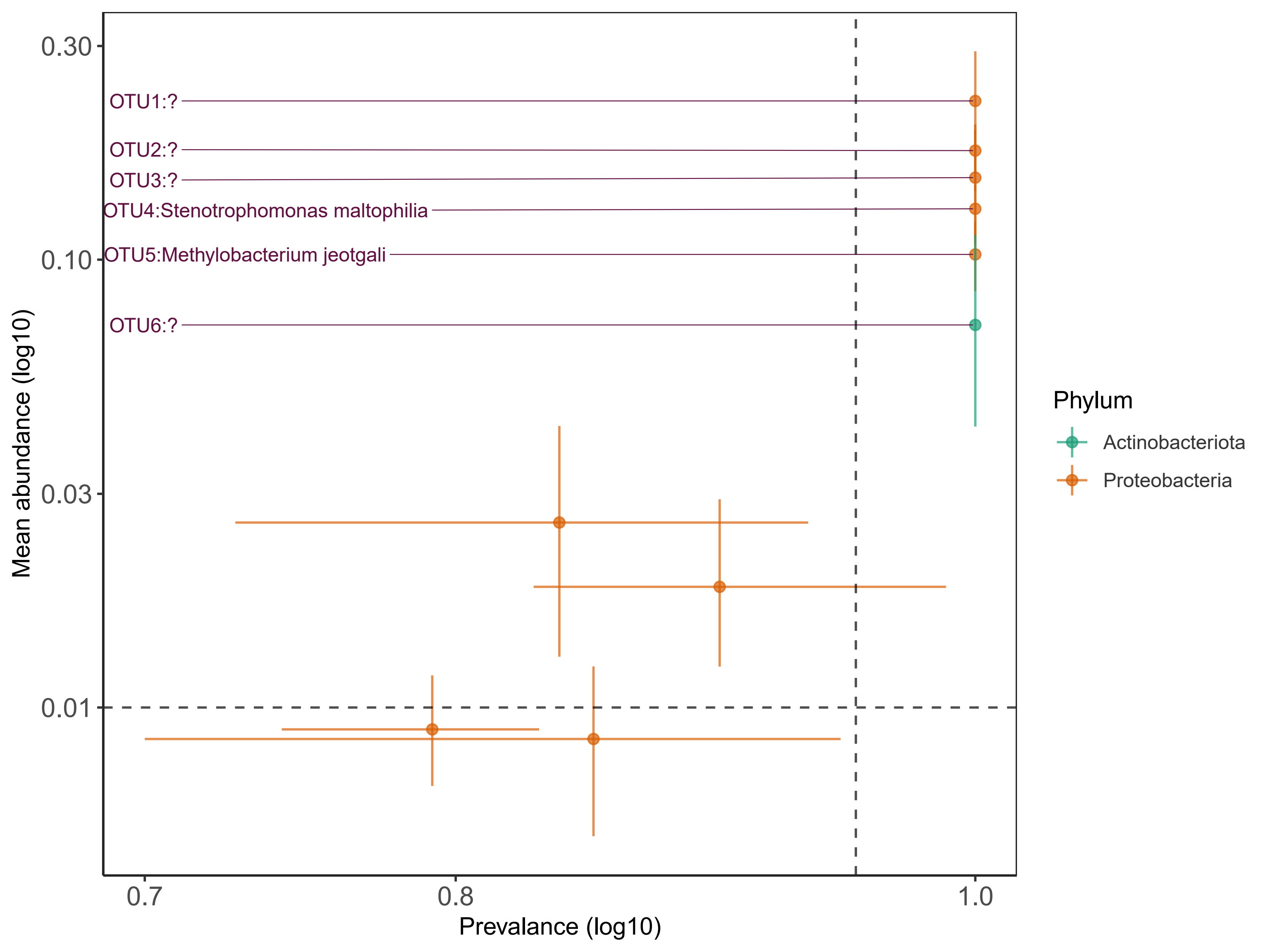

### Supp_Fig5_updated.pdf

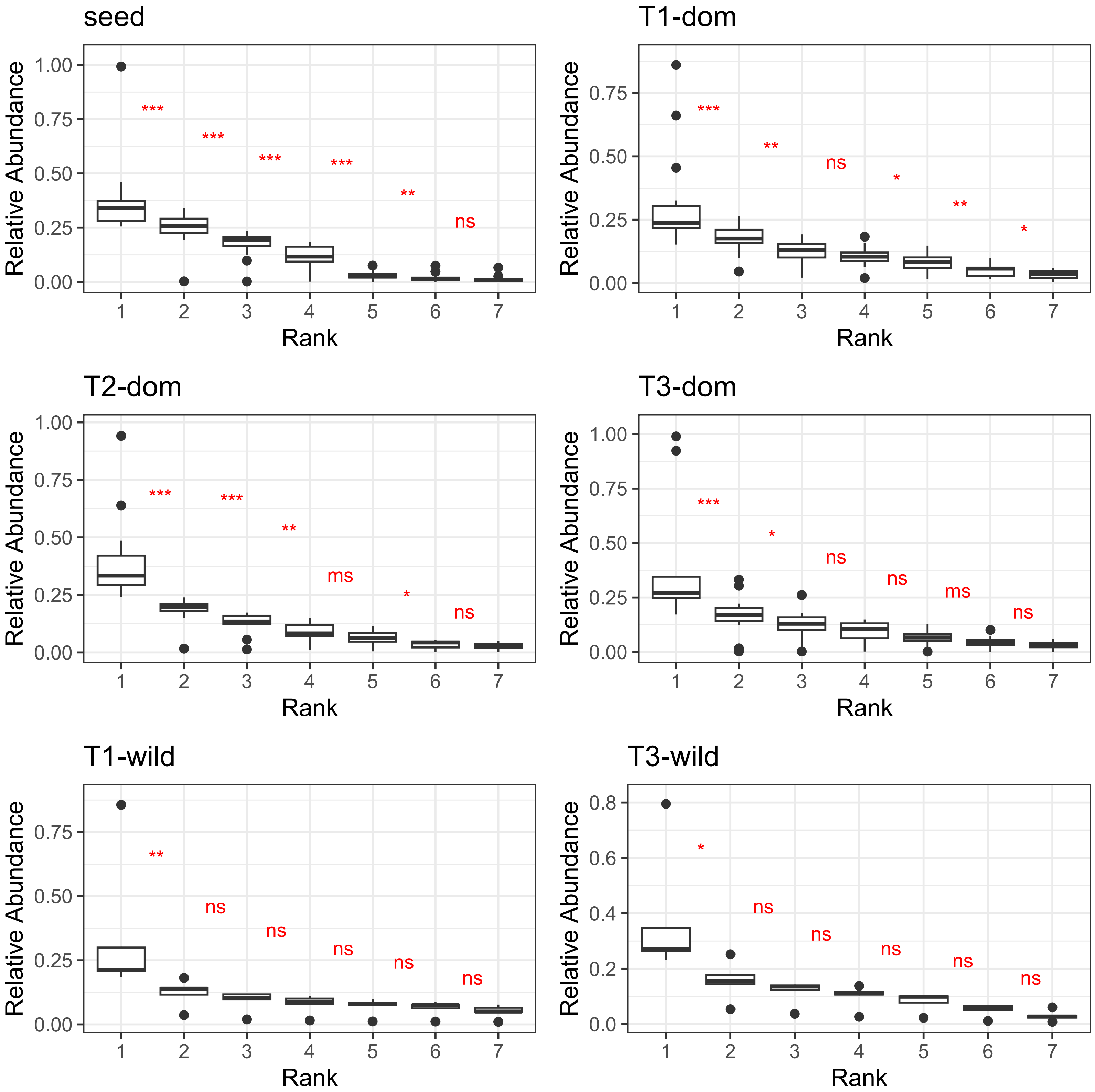

### Supp_Fig7_updated.pdf

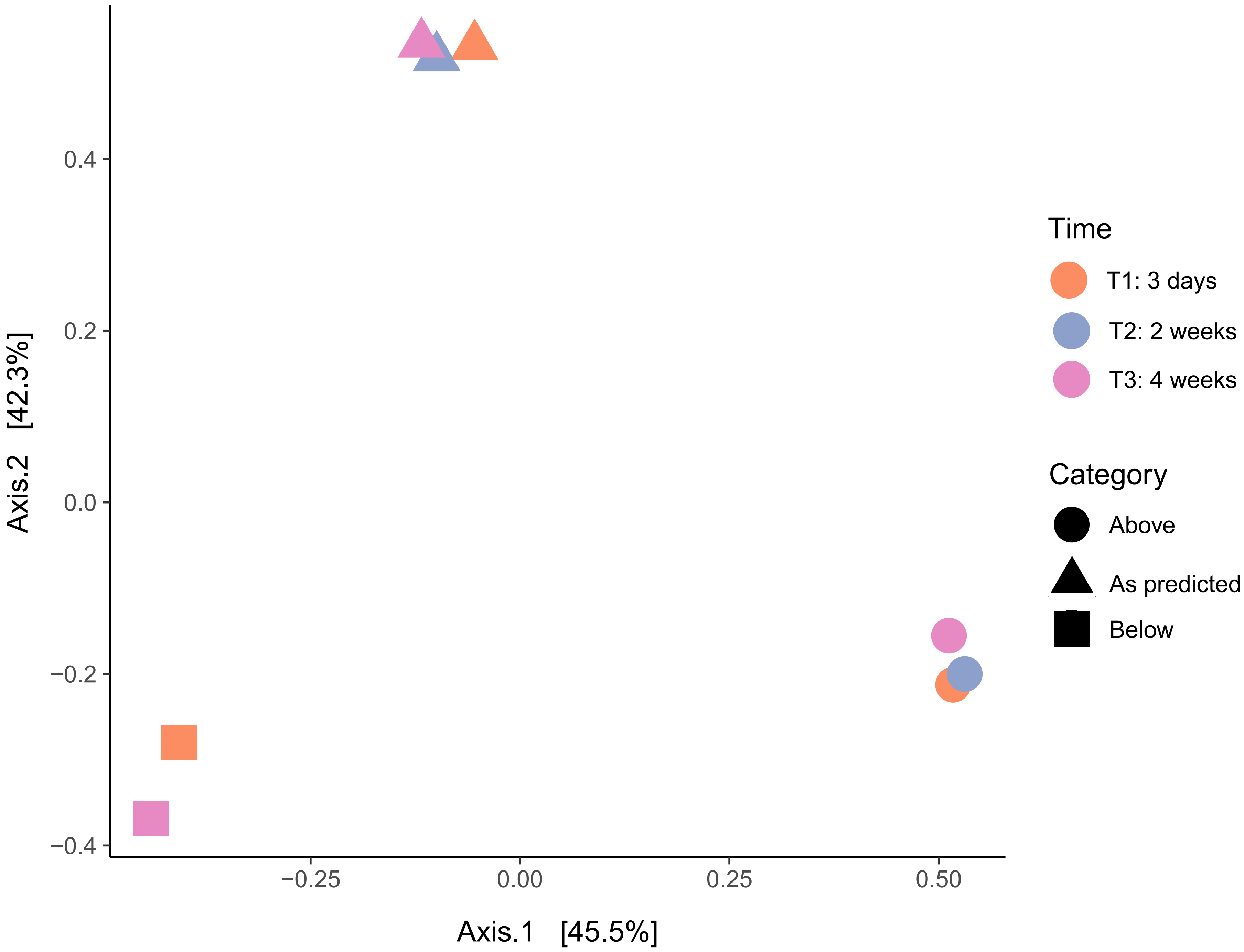

### SuppFig1_updated.pdf

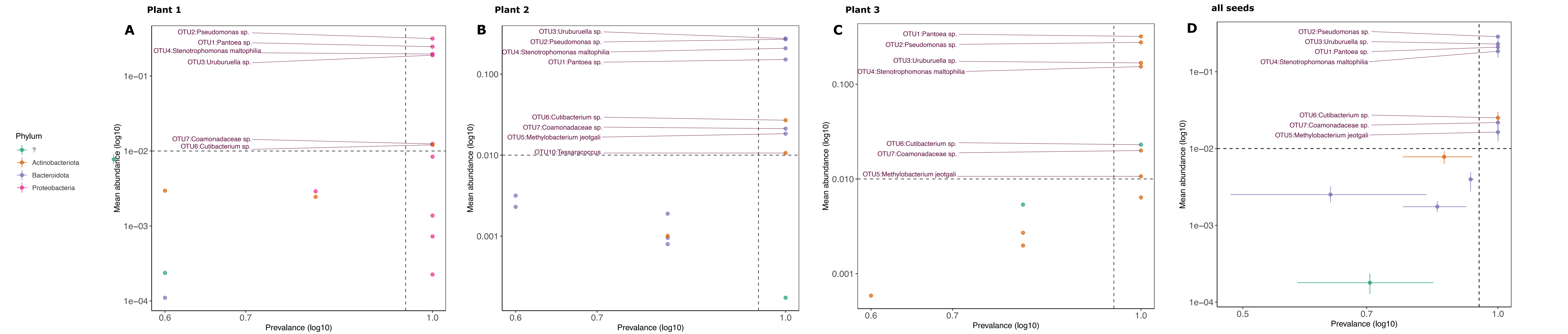

### SuppFig6_updated.pdf

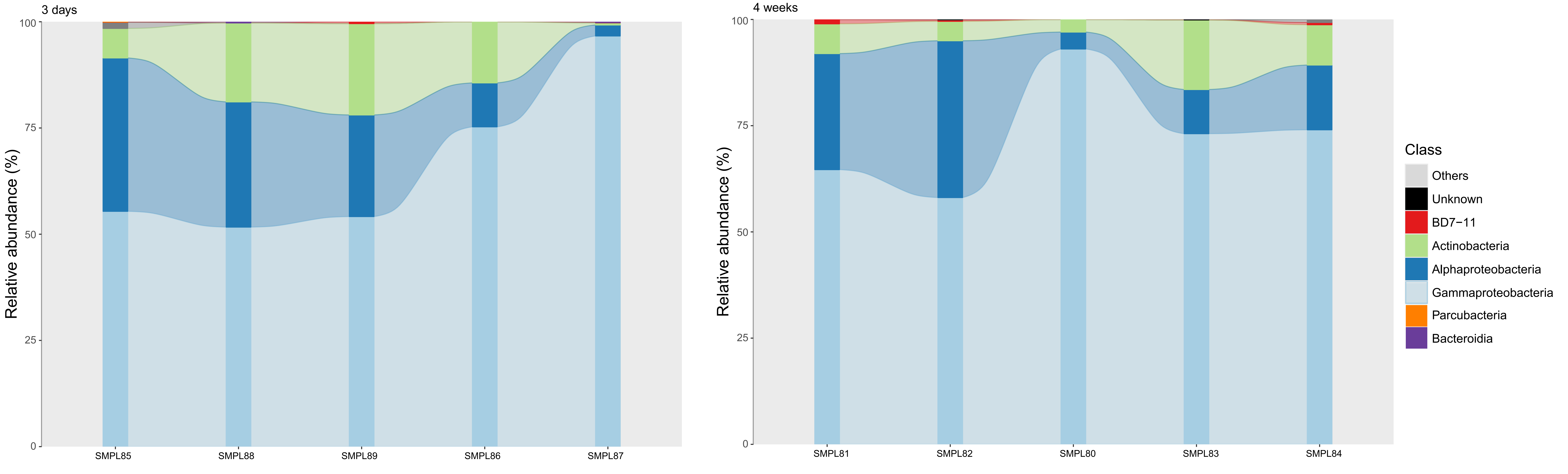
